# Regulatory network-based imputation of dropouts in single-cell RNA sequencing data

**DOI:** 10.1101/611517

**Authors:** Ana Carolina Leote, Xiaohui Wu, Andreas Beyer

## Abstract

Single-cell RNA sequencing (scRNA-seq) methods are typically unable to quantify the expression levels of all genes in a cell, creating a need for the computational prediction of missing values (‘dropout imputation’). Most existing dropout imputation methods are limited in the sense that they exclusively use the scRNA-seq dataset at hand and do not exploit external gene-gene relationship information. Further, it is unknown if all genes equally benefit from imputation or which imputation method works best for a given gene.

Here, we show that a transcriptional regulatory network learned from external, independent gene expression data improves dropout imputation. Using a variety of human scRNA-seq datasets we demonstrate that our network-based approach outperforms published state-of-the-art methods. The network-based approach performs particularly well for lowly expressed genes, including cell-type-specific transcriptional regulators. Further, the cell-to-cell variation of 12.6% to 48.2% of the genes could not be adequately imputed by any of the methods that we tested. In those cases gene expression levels were best predicted by the mean expression across all cells, i.e. assuming no measurable expression variation between cells. These findings suggest that different imputation methods are optimal for different genes. We thus implemented an R-package called ADImpute (available via Bioconductor https://bioconductor.org/packages/release/bioc/html/ADImpute.html) that automatically determines the best imputation method for each gene in a dataset.

Our work represents a paradigm shift by demonstrating that there is no single best imputation method. Instead, we propose that imputation should maximally exploit external information and be adapted to gene-specific features, such as expression level and expression variation across cells.

**Author summary:** Single-cell RNA-sequencing (scRNA-seq) allows for gene expression to be quantified in individual cells and thus plays a critical role in revealing differences between cells within tissues and characterizing them in healthy and pathological conditions. Because scRNA-seq captures the RNA content of individual cells, lowly expressed genes, for which few RNA molecules are present in the cell, are easily missed. These events are called ‘dropouts’ and considerably hinder analysis of the resulting data. In this work, we propose to make use of gene-gene relationships, learnt from external and more complete datasets, to estimate the true expression of genes that could not be quantified in a given cell. We show that this approach generally outperforms previously published methods, but also that different genes are better estimated with different methods. To allow the community to use our proposed method and combine it with existing ones, we created the R package ADImpute, available through Bioconductor.

## Introduction

Single-cell RNA sequencing (scRNA-seq) has become a routine method, revolutionizing our understanding of biological processes as diverse as tumor evolution, embryonic development, and ageing. However, current technologies still suffer from the problem that large numbers of genes remain undetected in single cells, although they actually are expressed (dropout events). Although dropouts are enriched among lowly expressed genes, relatively highly expressed genes can be affected as well. Of course, the dropout rate is also dependent on the sampling depth, i.e. the number of reads or transcript molecules (UMIs) quantified in a given cell. Imputing dropouts is necessary for fully resolving the molecular state of the given cell at the time of the measurement. In particular, genes with regulatory functions -e.g. transcription factors, kinases, regulatory ncRNAs -are typically lowly expressed and hence particularly prone to be missed in scRNA-seq experiments. This poses problems for the interpretation of the experiments if one aims at understanding the regulatory processes responsible for the transcriptional makeup of the given cell.

A range of computational methods have been developed to impute dropouts using the expression levels of detected genes. The underlying (explicit or implicit) assumption is often that detected and undetected genes are subject to the same regulatory processes, and hence detected genes can serve as a kind of ‘fingerprint’ of the state at which the cell was at the time of lysis. Several popular methods are based on some type of grouping (clustering) of cells based on the similarity of their expression patterns. Missing values are then imputed as a (weighted) average across those similar cells where the respective gene was detected(1–4). For example, the MAGIC algorithm(1) creates a network of cells by linking cells with similar gene expression signatures. Missing values are subsequently imputed by computing an average over linked cells, where cells get weighted based on how similar or dissimilar their expression signatures are compared to the target cell. DrImpute(3) and scImpute(2) have further developed this notion and have been shown to outperform MAGIC in recent comparisons(5). These methods rest on two important assumptions: (1) the global expression pattern of a cell (i.e. across the subset of detected genes) is predictive for all genes; (2) the (weighted) average of co-clustering (i.e. similar) cells is a good estimator of the missing value. The first assumption is violated if the expression of a dropout gene is driven by only a small subset of genes and hence the global expression pattern does not accurately reflect the state of the relevant sub-network. Any global similarity measure of the whole transcriptome will be dominated by the majority of genes(5). The second assumption is violated if the data is scarce, i.e. when either only few similar cells were measured or if the particular gene was detected in only a small subset of cells. In that case the average is computed across a relatively small number of observations and hence unstable.

Methods like SAVER(6) or G2S3(7) employ a different strategy that can overcome some of these limitations. Instead of using the whole transcriptome of a cell to predict the expression level of a given gene, these methods learn gene-gene relationships from the dataset and use only the specific subset of genes that are expected to be predictive for the particular gene at hand. For example, SAVER learns gene-gene relationships using a penalized regression model, whereas G2S3 optimizes a sparse gene graph.

SCRABBLE is different compared to all of the other methods mentioned above, because it can use bulk sequencing data to assist in the imputation. SCRABBLE combines a de-noising step with a moderated imputation moving the sample means towards the observed (bulk-derived) mean expression values.

Here, we compare published approaches that are representative for current state-of-the-art methods to two fundamentally different approaches. The first is a very simple baseline method that we use as a reference approach: we estimate missing values as the average of the expression level of the given gene across all cells in the dataset where the respective gene was detected. Initially intended to serve just as a reference for minimal expected performance, this sample-wide averaging approach turned out to perform surprisingly well and in many instances even better than state-of-the-art methods. The simple explanation is that estimating the average using all cells is a much more robust estimator of the true mean than using only a small set of similar cells, especially when the gene was detected in only few cells and/or if the gene’s expression does not vary much across cells.

The second new approach avoids using a global similarity measure comparing entire transcriptomes. Instead, similar to SAVER or G2S3 it rests on the notion that genes are part of regulatory networks and only a small set of correlated or functionally associated genes should be used to predict the state of undetected genes. However, unlike other methods, we propose to use transcriptional regulatory networks trained on independent (bulk seq) data to rigorously quantify the transcriptional relationships between genes. Missing values are then imputed using the expression states of linked genes in the transcriptional regulatory network and exploiting the known quantitative relationships between genes. This approach allows imputing missing states of genes even in cases where the respective gene was not detected in any cell or in only extremely few cells. This second new approach rests on the assumption that the network describes the true regulatory relationships in the cells at hand with sufficient accuracy. Here, we show that this is indeed the case and that combining the two new approaches with published state-of-the-art methods drastically improves the imputation of scRNA-seq dropouts. Importantly, the performance of an imputation method is dependent on the ‘character’ of a gene (e.g. its expression level or the variability of expression between cells). Hence, we implemented an R-package (Adaptive Dropout Imputer, or ADImpute) that determines the best imputation method for each gene through a cross-validation approach.

## Results

### Imputing dropouts using a transcriptional regulatory network

In order to understand whether the inclusion of external gene regulatory information allows for more accurate scRNA-seq dropout imputation, we derived a regulatory network from bulk gene expression data in 1,376 cancer cell lines with known karyotypes. While the expression levels of genes in this data will be cell type-specific, the relationships between genes (e.g. concerted up-regulation of a transcription factor and its targets) are frequently conserved across cell types, allowing us to pool the data together to learn a generic gene regulatory network. For this purpose, we modelled the change (compared to average across all samples) of each gene as a function of its own copy number state and changes in predictive genes:

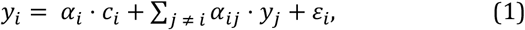

where *y*_*i*_ is the expression deviation (log fold change) of gene *i* from the global average, *c*_*i*_ is the known (measured) copy number state of gene *i, α* the vector of regression coefficients, *y*_*j*_ the observed change in expression of gene *j* and *ε*_*i*_ the i.i.d. error of the model. To estimate a set of predictive genes *j*, we made use of LASSO regression(8), which penalizes the L1 norm of the regression coefficients to determine a sparse solution. LASSO was combined with stability selection(9) to further restrict the set of predictive genes to stable variables and to control the false discovery rate (Methods). This approach ensures that the algorithm only selects gene-gene relationships that are invariant across most or all training data. Thus, interactions that would be specific to a single cell type will be excluded from the model. Using the training data, models were fit for 24,641 genes, including 3,696 non-coding genes. The copy number state was only used during the training of the model, since copy number alterations are frequent in cancer and can influence the expression of affected genes. If copy number states are known, they can of course also be used during the dropout imputation phase. Using cell line data for the model training has the advantage that the within-sample heterogeneity is much smaller than in tissue-based samples(10). However, in order to evaluate the general applicability of the model across a wide range of conditions, we validated its predictive power on a diverse set of tissue-based bulk-seq expression datasets from the The Cancer Genome Atlas (4,548 samples from 13 different cohorts; see Methods and Supplementary Fig. 1) and the Genotype-Tissue Expression (17,382 samples from 30 different healthy tissues; see Methods and Supplementary Fig. 2).

**Figure 1:**
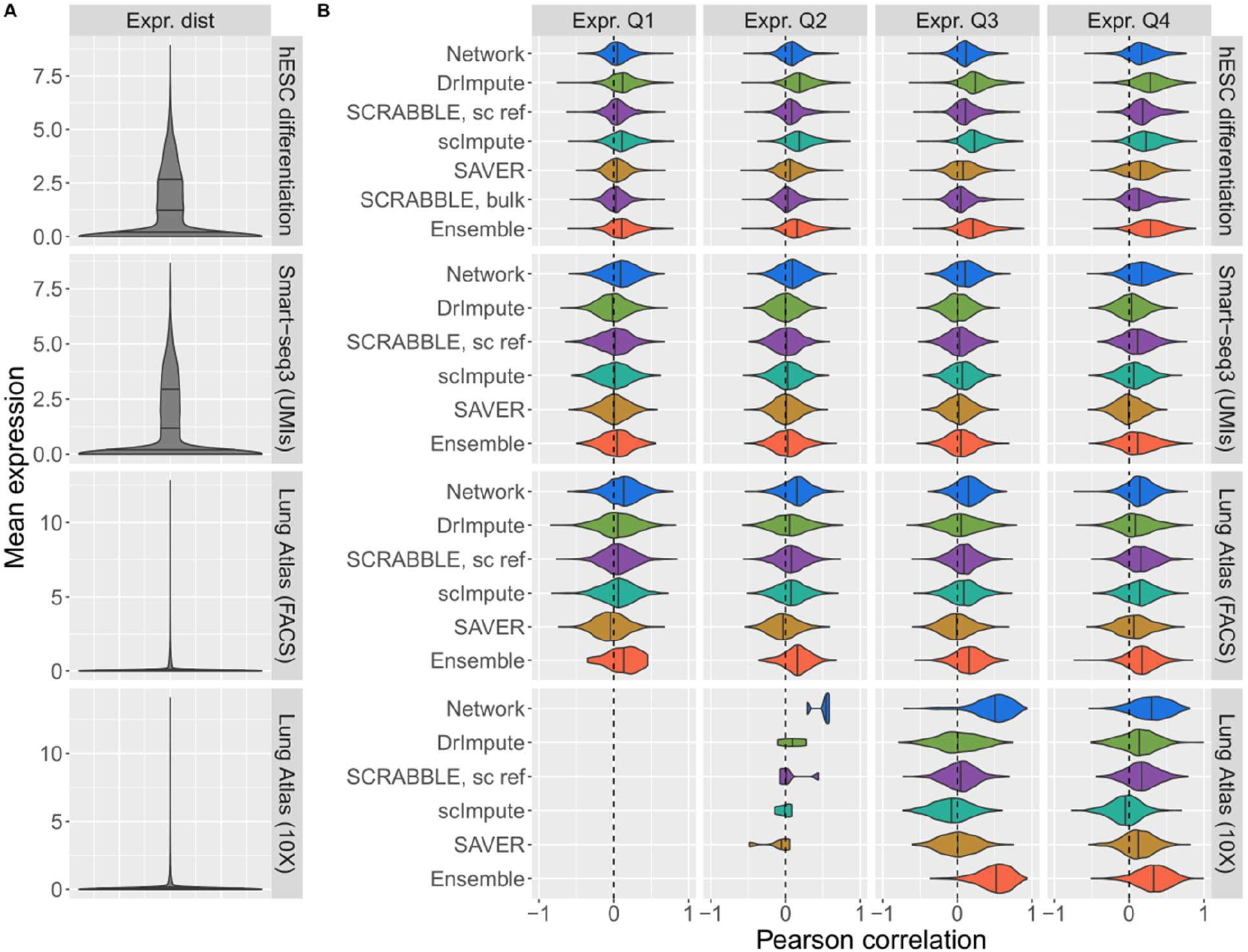
Imputation performance of Network (blue), DrImpute (green), SCRABBLE (purple), scImpute (turquoise), SAVER (yellow) and Ensemble (orange). **A)** Distribution of average expression levels (per gene) in each dataset. Quartiles are represented by vertical lines. B) Pearson correlation coefficient, for each gene, between the imputation by the specified method and the original values before masking. Only values that could be imputed by all methods (non-zero imputation) were considered. Correlations per gene across cells were computed for all genes for which at least 10 imputation values were available for analysis. Expression quartiles were determined for each dataset separately, on the masked data.

**Figure 2:**
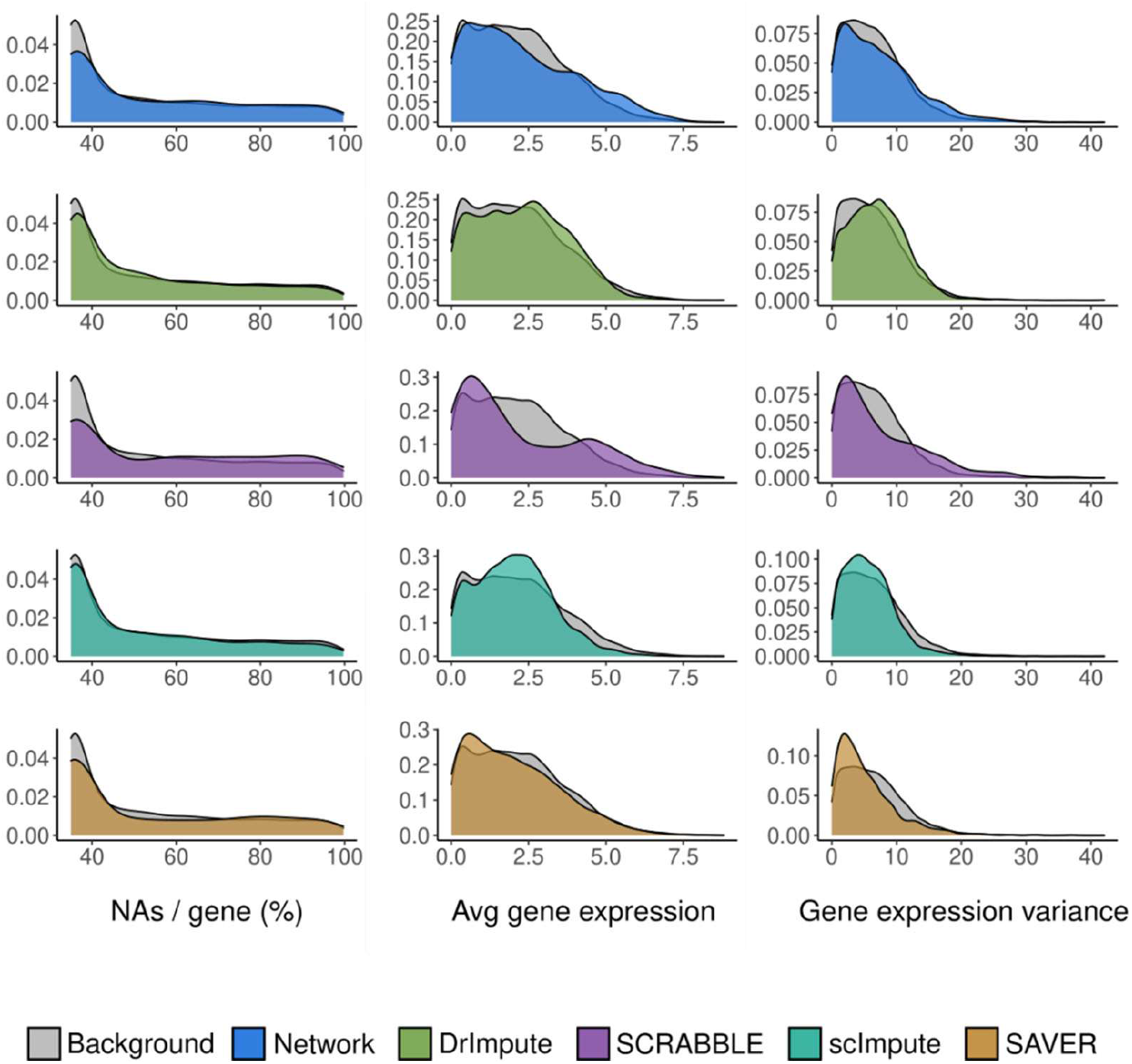
Characterization of the genes best predicted by Network (blue), DrImpute (green), SCRABBLE (bulk data as reference; purple), scImpute(turquoise) and SAVER (yellow) in the hESC differentiation dataset. Distribution of missing values per gene, average expression levels and variance of the genes best predicted by Baseline, scImpute and Network methods, compared against all tested genes (background). Average gene expression is shown as log_2_-transformed normalized expression.

Such a model allows us to estimate the expression of a gene that is not quantified in a given cell based on the expression of its predictors in the same cell. Here, the difficulty lies in the fact that imputed dropout genes might themselves be predictors for other dropout genes, i.e. the imputed expression of one gene might depend on the imputed expression of another gene. In order to derive the imputation scheme based on the model from equation (1), we revert to an algebraic expression of the problem,

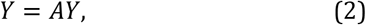

where *A* is the adjacency matrix of the transcriptional network, with its entries *a*_*ij*_ being fitted using the regression approach described above, and *Y* is the vector of gene expression deviations from the mean across all cells in a given cell. In the current implementation we assume no copy number changes and hence, we exclude the *c*_*i*_ term from equation (1). Like in equation (1), we omit the intercept since we are predicting the deviation from the mean. Subsequently, imputed values are re-centered using those means to shift imputed values back to the original scale (see Methods). Further note that we drop the error term *ε* from equation (1), because this is now a prediction task (and not a regression). Here, we exclusively aim to predict dropout values, and (unlike SAVER) our goal is not to improve measured gene expression values. Hence, measured values remain unchanged. It is therefore convenient to further split *Y*into two sub-vectors *Y*^*m*^and *Y*^*n*^, representing the measured and non-measured expression levels, respectively. Likewise, *A* is reduced to the rows corresponding to non-measured expression levels and split into *A*^*m*^ (dimensionality |*n*| *x* |*m*|) and *A*^*n*^ (dimensionality |*n*| × |*n*|), accounting for the contributions of measured and non-measured genes, respectively. The imputation problem is then reduced to:

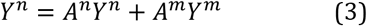

As *Y*^*m*^ is known (measured) and will not be updated by our imputation procedure, the last term can be condensed in a fixed contribution, *F = A*^*m*^*Y*^*m*^, accounting for measured predictors:

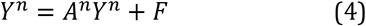

The solution *Y*^*n*^ for this problem is given by:

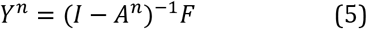

The matrix *(I − A*^*n*^) may not be invertible, or if it is invertible, the inverse may be unstable. Therefore, we computed the pseudoinverse *(I − A*^*n*^)^*+*^ using the Moore-Penrose inversion. Computing this pseudoinverse for every cell is a computationally expensive operation. Thus, we implemented an additional algorithm finding a solution in an iterative manner (Methods). Although this iterative second approach is not guaranteed to converge, it did work well in practice (see Supplementary Fig. 3, Methods). While our R-package implements both approaches, subsequent results are based on the iterative procedure.

**Figure 3:**
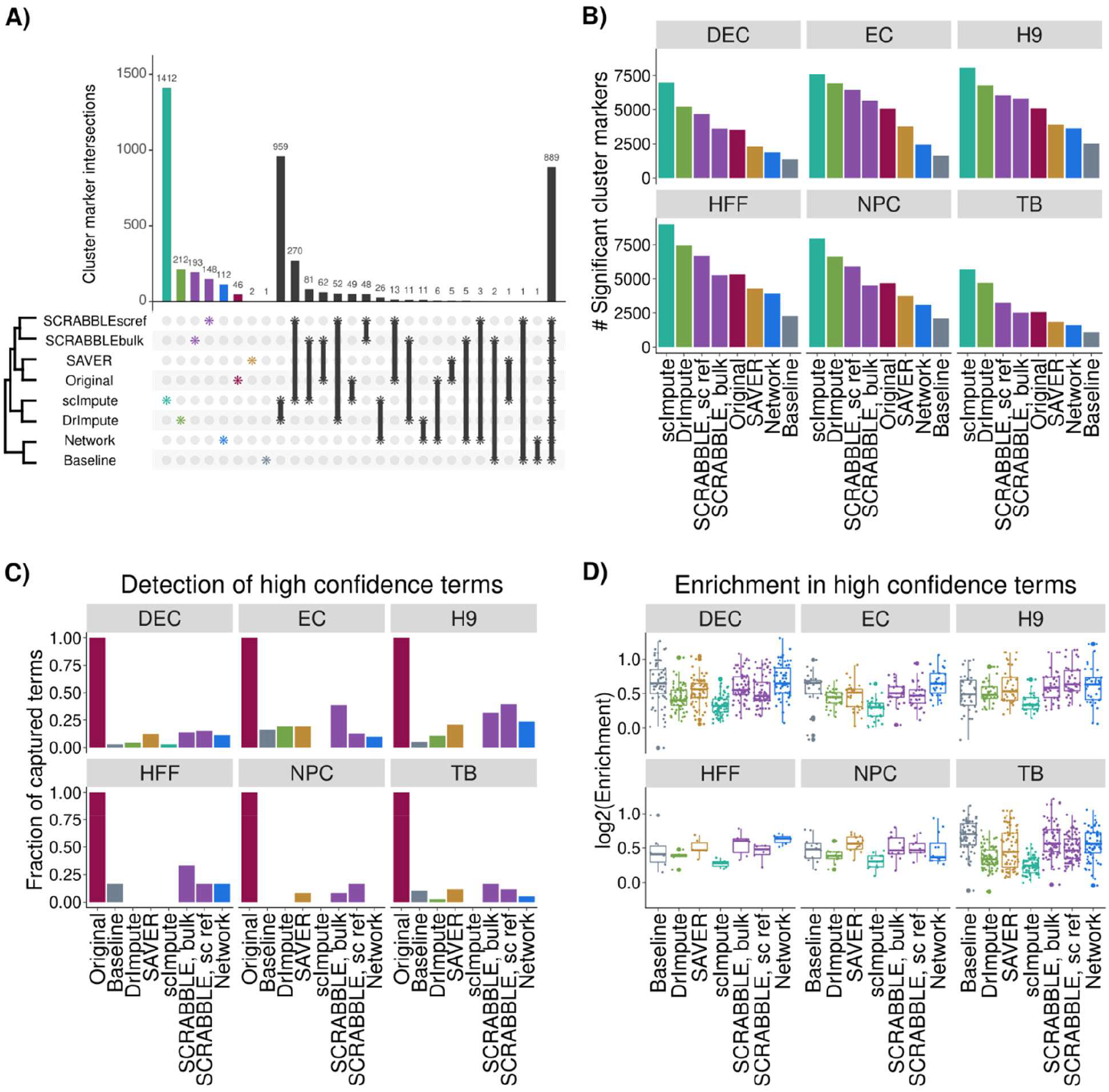
Detection of cell type-specific markers before and after imputation. **A)** Overlap between significant (FDR < 0.05, |log_2_FC| > 0.25) definitive endoderm cell markers detected with no dropout imputation (Original) and using the tested imputation methods. **B)** Number of significant cell type markers detected with no dropout imputation and using the tested imputation methods. **C)** and **D)** fraction of captured high confidence terms, defined as significantly enriched (p.value < 0.001 and log_2_Enrichment > 0.5, Methods) GO biological process terms among the cluster markers detected without imputation. **C)** Fraction of high confidence terms detected as significantly enriched (p.value < 0.001 and log_2_Enrichment > 0.5) among the cluster markers detected with each imputation method. **D)** log_2_-enrichment of all high confidence terms among the cluster markers detected with each imputation method. DEC: definitive endoderm cells; EC: endothelial cells; H9: undifferentiated human embryonic stem cells; HFF: human foreskin fibroblasts; NPC: neural progenitor cells; TB: trophoblast-like cells.

### Transcriptional regulatory network information improves scRNA-seq dropout imputation

To assess the performance of our network-based imputation method and compare it to that of previously published methods, we considered seven different single-cell RNA-sequencing datasets (11–15), covering a wide range of sequencing techniques (Smart-seq versions 1, 2 and 3 and droplet-based method 10X) and biological contexts (healthy tissue, cancer, stem cell differentiation and HEK cells). A summary of the dataset characteristics, including number of cells and average number of quantified genes per cell, is provided in Supplementary Table 1. It was important to include a range of different healthy cell types in the evaluation, because the transcriptional regulatory network was trained on cancer cell line data. Thus, by including data from non-cancerous tissues, we could evaluate possible restrictions induced by the model training data.

**Table 1:**
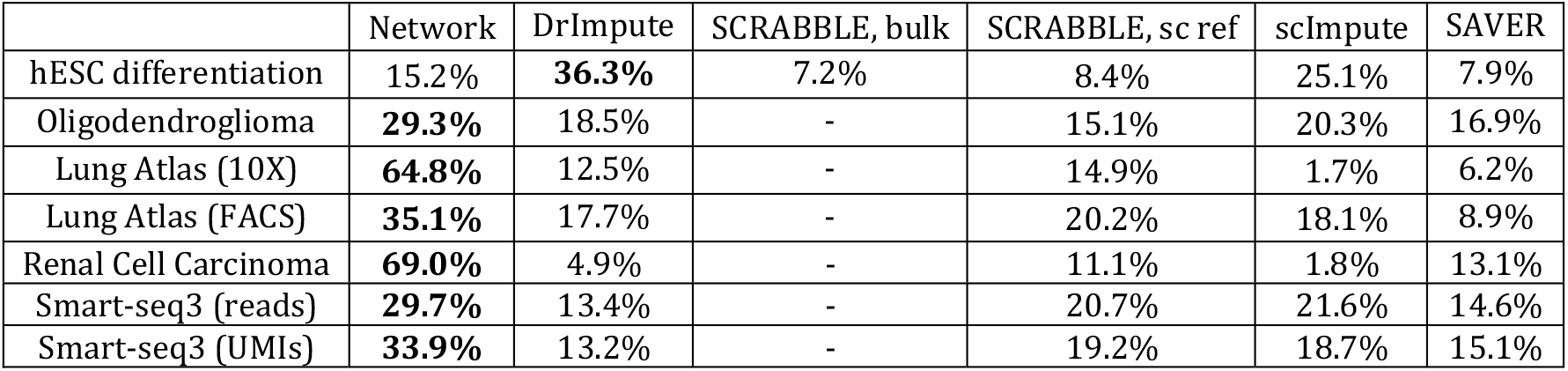
Percentage of genes best imputed by each method (highest Pearson correlation coefficient) in the seven test datasets restricted to values that could be imputed by all individual methods.

In order to quantify the performance of both proposed and previously published imputation methods, we randomly set a fraction of the quantified values in the test data to zero according to two different schemes (Methods) and stored the original values for later comparison with the imputed values. Imputation was then performed on the masked dataset using our network-based approach, DrImpute (3), SAVER (6), scImpute (2) and SCRABBLE (16). Those methods were chosen since they were shown to be among the top-performing state-of-the-art dropout imputation methods (17). For masked entries imputed by all tested methods, the quality of imputation was assessed for each gene, using two approaches: computing the correlation between observed and imputed values for each gene, and computing the Mean Squared Error of imputation (Methods).

According to the correlation quality measure our network-based approach (called ‘Network’) mostly outperformed other methods (Figure 1). We quantified the percentage of genes in the transcriptome of each dataset best imputed by each method (highest correlation) and verified that in six out of our seven test datasets, the network-based approach resulted in the highest performance for most genes (Table 1). Additionally, we observed that Network was less affected by low average expression levels compared to all other imputation methods (Figure 1, Supplementary Fig. 4, expression quartiles Q1 and Q2). This was expected since our network-based approach relies on information external to the dataset for dropout imputation, while other methods require sufficient observations of a gene to learn its expression characteristics from the single cell data itself. This result is in line with, for instance, a previous observation that scImpute is sensitive to missing information about genes across cells (17). SCRABBLE is able to incorporate the average expression in matched bulk RNA-seq data to aid imputation, effectively taking advantage of external information like Network. Although this information is not available for most scRNA-seq datasets, it was available for the hESC dataset, prompting us to use it in an additional SCRABBLE test (Figure 1, Supplementary Figs. 4-5). We observed that SCRABBLE’s performance was not improved when incorporating this additional information.

**Figure 4:**
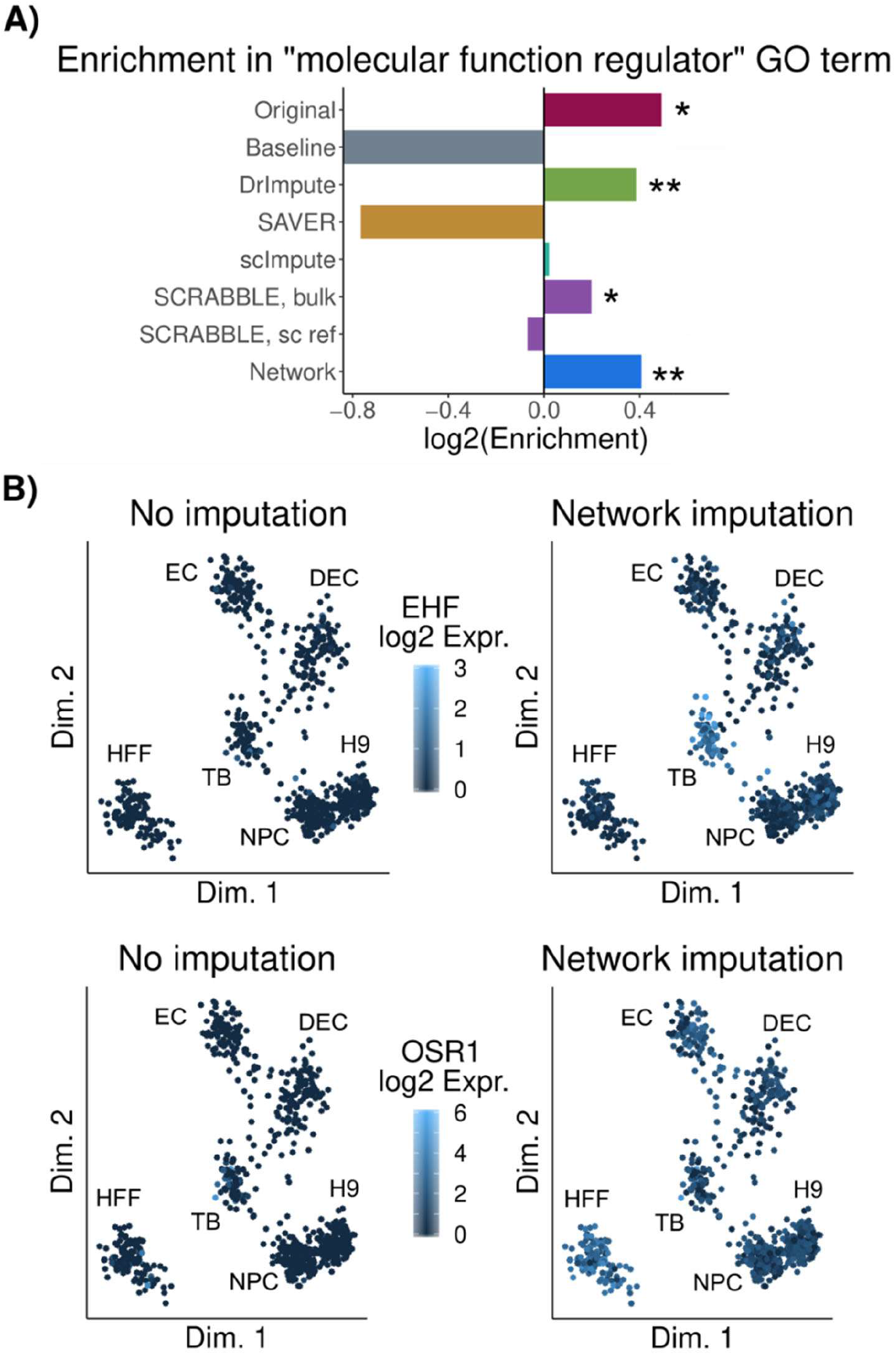
Detection of cell type-specific transcription factors is improved upon network-based imputation. **A)** Enrichment score in GO term “molecular function regulator” among the genes uniquely detected after each imputation approach. **: p-val < 0.01; *: p-val < 0.05. B) Projection of cells onto a low dimension representation of the data before imputation, using ZINB-WaVe(24). Color represents normalized expression levels of EHF (top) and OSR1 (bottom) before and after Network-based imputation. DEC: definitive endoderm cells; EC: endothelial cells; H9: undifferentiated human embryonic stem cells; HFF: human foreskin fibroblasts; NPC: neural progenitor cells; TB: trophoblast-like cells.

As the network-based approach uses information regarding other genes contained in the same cell, we hypothesized its accuracy might be more affected by increasingly sparse information per cell when compared to other methods. However, the relative performance differences between methods were largely invariant to the number of missing genes per cell (Supplementary Fig. 7), suggesting that other methods also suffer from the scarcity of information for cells with low sampling efficiency. The sensitivity of the imputation to the proportions of missing values was dependent on a multitude of factors, including the quality of the data, the specific gene(s) expressed in the cell type(s) at hand, the number of missing values, variability of expression/cell type heterogeneity within the study and possibly many others. Further, the network-based method performed well over a range of different cell types and showed decreased performance upon randomization of the transcriptional network (Methods, Supplementary Fig. 8). Thus, the diversity of cell lines used in the training data seemed to capture a large fraction of all possible regulatory relationships in the human transcriptome.

We additionally evaluated imputation performance using the mean squared error (MSE) between original and imputed values. In this analysis, we included a ‘Baseline’ method, which imputes dropouts with the average expression value of the gene across all cells, as a reference. Using the MSE as an alternative metric of performance, we also observed that Network outperformed previously published methods across all tested datasets and expression quartiles (Supplementary Fig. 6). However, the performance of the Baseline method (Supplementary Fig. 6, grey violin), which does not account for any expression variation between cells, was surprising to us. While SAVER showed a poor performance in comparison to all other methods (also described in (17)), it should be noted that this method aims to estimate the true value for all genes, not only for the dropout genes. This results in a change of both dropout and quantified values, which explains the good correlation performance but relatively high MSE.

Taken together, these results indicate that Network often – but not always -leads to more accurate (Supplementary Fig. 6) imputations than state-of-the-art imputation methods, while preserving variation across cells as captured by the correlation analysis (Figure 1, Supplementary Fig. 4). However, our analysis also uncovered that the advantage of using specific methods varies between datasets and expression quartiles, suggesting that there is no universally best performing method that outperforms the others in all cases. This motivated us to develop an ensemble approach, where we determine in a cross-validation scheme the best imputation method for each gene in the dataset at hand. We tested its performance (Figure 1, Supplementary Figs. 4 and 6) and observed that the ensemble method tends to approach the performance of the best performing method.

### Gene features determine the best performing imputation method

To better understand what gene features drive the performance differences between methods, we characterized the genes best imputed by each of the methods. We determined, for each gene in each test dataset, the method resulting in the highest correlation between imputed and original values (Table 1). The genes best imputed by each method were then compared against a background including all genes for which all methods were able to perform imputations (Fig. 2 and Supplementary Fig. 10). As expected, methods relying on the similarity of cellular transcriptomes (scImpute, DrImpute) performed best for more frequently detected genes (lower percentage of NAs per gene across cells). Conversely, SAVER, SCRABBLE and Network outperformed the other methods especially on genes with many missing values. SAVER and Network are model-based methods, not relying on the comparison of entire transcriptomes between cells. SCRABBLE and Network are methods using external information for the imputation. Based on the above results we conclude that both aspects (model-based imputation and using external information) are advantageous for the imputation of rarely detected genes. We also determined the best performing method for each gene based on the MSE instead of correlations. We observed that, as expected, the Baseline method performed best for genes with low expression levels and variance (Supplementary Fig. 11).

### Network-based imputation uncovers cluster markers and regulators

A popular application of scRNA-seq is the identification of discrete sub-populations of cells in a sample in order to, for example, identify new cell types. The clustering of cells and the visual 2D representation of single-cell data is affected by the choice of the dropout imputation method^13^. Therefore, we assessed the impact of dropout imputation on data visualization using Uniform Manifold Approximation and Projection (UMAP)(18) on the hESC data before and after imputation by all methods. The hESC dataset was particularly suitable in this case, because it was of high quality and it consisted of six well-annotated cell types. This analysis confirmed that the choice of the imputation method impacts on the grouping/clustering of cells (Supplementary Fig. 12). Application of other dimensionality reduction techniques (t-SNE, ZINBWaVe) showed varying results depending on the chosen method, suggesting that visual clustering upon dimensionality reduction is an inconclusive criterion for evaluating dropout imputation (Supplementary Fig. 13).

We next asked to what extent the detection of cluster markers would be affected by the choice of the imputation method. Thus, we applied Seurat(19) to the hESC differentiation dataset, which was composed of a well-defined set of distinct cell types, before and after imputation. We then defined genes that were significantly differentially expressed between one cluster and all the others as cluster markers (Methods). We observed a strong overlap between markers detected before and after applying the tested imputation methods (Fig. 3A, rightmost bar, Supplementary Fig. 14), suggesting a common core of detected cluster markers across methods. Additionally, the numbers of significant markers detected after Network and Baseline imputations were lower than for other imputation methods (Fig. 3B). Imputation with scImpute and, to a smaller extent, with DrImpute, led to the highest number of significant markers (Fig. 3B). We hypothesized that many of these marker genes may result from artefactual clustering of cells. In order to test that notion we first determined all GO biological process terms that were enriched in the respective cell clusters without any dropout imputation. We termed them ‘high confidence GO terms’ since they are independent of the choice of the imputation method. It turned out that scImpute and DrImpute had the weakest enrichments in high confidence GO biological process terms (Fig. 3C-D; Methods; Supplementary Table 4), suggesting that the extra markers found upon applying scImpute and DrImpute contained many false positives, which diluted biological signals. Conversely, Network and SCRABBLE led to the strongest enrichments in high confidence GO biological process terms (Fig. 3C-D).

Genes with regulatory functions are particularly important for understanding and explaining the transcriptional state of a cell. However, since genes with regulatory functions are often lowly expressed(20), they are frequently subject to dropouts. Since our analysis had shown that the network-based approach is especially helpful for lowly expressed genes (Fig. 2), we hypothesized that the imputation of transcript levels of regulatory genes would be particularly improved. In order to test this hypothesis, we further characterized those cluster markers that were exclusively detected using the network-based method. Indeed, we observed regulatory genes to be enriched among those markers (Fig. 4A). Among these markers exclusively detected upon network-based imputation, the transcription factor EHF was the second most significant trophoblast-specific. EHF is a known epithelium-specific transcription factor that has been described to control epithelial differentiation(21) and to be expressed in trophoblasts(22), even though at very low levels (EHF expression found among the first quintile of bulk TB RNA-seq data from the same authors). While EHF transcripts were not well captured in TB single-cell RNA-seq data (only quantified in 39 out of 775 TB cells), a trophoblast-specific expression pattern was recovered after network-based imputation (Fig. 4B, upper panel), but not with any of the other tested imputation methods (Supplementary Fig. 15). Similarly, OSR1 has been described as a relevant fibroblast-specific transcription factor(23) which failed to be detected without imputation. Imputing with Network lead to the strongest fibroblast-specific expression pattern of OSR1 (Fig. 4B, lower panel), (Supplementary Fig. 15). Interestingly, TWIST2 and PRRX1, described by Tomaru *et al*.(23) to interact with OSR1, also showed fibroblast-specific expression (Supplementary Fig. 16). Taken together, these results suggest that imputation based on transcriptional regulatory networks can recover the expression levels of relevant, lowly expressed regulators affected by dropouts.

## Discussion

This work has led to the following key findings: (i) a model-based approach using external data is particularly powerful for the imputation of rarely detected genes; (ii) based on the MSE criterion the expression of surprisingly many genes is best predicted by simply using their average expression across cells (‘Baseline’); (iii) not all genes are equally well predicted by a single imputation approach; instead, one should adapt the imputation method to the specific gene in a given dataset. In addition, our work confirmed earlier findings, such as the artifactual clustering resulting from some imputation methods.

The consideration of external gene co-expression information for the dropout imputation substantially improved the performance in many cases, especially for lowly expressed genes. Since genes with regulatory functions are often lowly expressed (20), imputation of those genes might be critical for explaining expression variation between cells. Of note, cell type-specific regulatory genes were successfully imputed using information from our global gene co-expression network (Fig. 4). This observation, together with the demonstrated predictive capacity of our network across cancer and healthy human data from a wide range of tissues, highlights the transferability of the gene-gene relationships learnt by our network.

A potential limitation of our approach is that a transcriptional network derived from bulk-seq data may not fully capture gene-gene relationships that are detectable from single cell data. For example, gene regulatory relationships that are specific to a small sub-population of cells in a bulk tissue may not be correctly captured, because the signal would be too weak. A second example would be genes regulated during the cell cycle. Bulk tissue is usually not synchronized, i.e. it consists of a mix of cells at different cell cycle stages, which may prevent the detection of those relationships. To some extent these limitations were alleviated by using cell line data rather than actual tissue data for training the network. Of course, the network that we used here is still imperfect. However, despite that imperfection it demonstrated the power of our approach. Using it was clearly advantageous over not using it in most dropout imputation tests, and we showed its predictive power across independent datasets from both cancer and healthy human tissue.

A surprising finding of our analysis is the fact that the sample-wide average expression (‘Baseline’) performs well for the imputation of many genes when using the MSE as a performance measure (Supplementary Table 3). As expected, genes whose expression levels were best imputed by this method were characterized by lower variance across cells and by remaining undetected in relatively many cells (Supplementary Fig.11). A potential problem of methods based on co-clustering cells is that the number of observations per cluster can get very small, which makes the estimation of the true mean more unstable. Thus, averaging across all cells is preferred when the gene was detected in only few cells and/or if the gene’s expression does not vary much across cells. Further, our findings imply that cell-to-cell expression variation of many genes is negligible or at least within the limits of technical measurement noise.

The third --and maybe most important --conclusion is that the best performing imputation method is gene-and dataset-dependent. That is, there is no single best performing method. If the number of observations is high (many cells with detected expression) and if the expression quantification is sufficiently good, scImpute and DrImpute outperformed other methods. Importantly, the technical quality of the quantification depends on the read counts, which in turn depends on sampling efficiency, gene expression, transcript length and mappability – i.e. multiple factors beyond expression. If however, gene expression is low and/or too imprecise, scImpute and DrImpute were outcompeted by other methods. This finding led us to conclude that a combination of imputation methods would be optimal. Hence, we developed an R-package that determines ‘on the fly’ for each gene the best performing imputation method by masking observed values (i.e. *via* cross validation). This approach has the benefit that it self-adapts to the specificities of the dataset at hand. For example, the network-based approach might perform well in cell types where the assumptions of the co-expression model are fulfilled, whereas it might fail (for the same gene) in other cell types, where these assumptions are not met. Hence, the optimal imputation approach is gene-and dataset-dependent. An adaptive method selection better handles such situations. Another benefit of this approach is that the cross validation imputation performance can be used as a quantitative guide on how ‘imputable’ a given gene is in a specific scRNA-seq dataset. We have therefore implemented and tested this approach (see Figure 1, Supplementary Figs. 4-5). The resulting R-package (called ADImpute) is open to the inclusion of future methods, includes scImpute’s estimation of dropout probability and it can be downloaded from https://bioconductor.org/packages/release/bioc/html/ADImpute.html.

We believe that this work presents a paradigm shift in the sense that we should no longer search for the single best imputation approach. Rather, the task for the future will be to find the best method for a particular combination of gene and experimental condition.

## Methods

### Pre-processing of cancer cell line data for transcriptional regulatory network inference

Entrez IDs and corresponding gene symbols were retrieved from the NCBI (https://www.ncbi.nlm.nih.gov/gene/?term=human%5Borgn%5D). Genome annotation was obtained from Ensembl (*Biomart*). Finally, genes of biotype in protein coding, ncRNA, snoRNA, scRNA, snRNA were used for network inference. For CCLE(25), 768 cell lines that were used in Seifert et al.^9^ were used. Raw CEL files were downloaded from https://portals.broadinstitute.org/ccle/ and processed using the R package RMA in combination with a BrainArray design file (HGU133Plus2_Hs_ENTREZG_21.0.0). Final expression values were in log_2_ scale. Expression levels and CNV data set from RNA-seq were downloaded from Klijn et al.(26). Before combining, each dataset is log_2_ transformed and scaled to (0,1) for all genes in each sample using R function scale. Then datasets were merged and the function ComBat from the sva R package(27) was used to remove batch effect of the data source. The final combined data set contains 24641 genes in 1443 cell lines. Finally, expression levels of genes were subtracted by the average expression level across all cell lines of the corresponding gene.

### Network inference based on stability selection

The network inference problem can be solved by inferring independent gene-speci?c sub-networks. We used the linear regression model from equation (1) to model the change in a target gene as dependent on the combination of the gene-specific CNA and changes in all other genes. Here the intercept is not included because the data is assumed to be centered. We used LASSO with stability selection(9) to find optimal model parameters *a*_*ij*_.

The R package *stabs* was employed to implement stability selection and the *glmnet* package was used to fit the generalized linear model. Two parameters regarding error bounds were set with the cutoff value being 0.6 and the per-family error rate being 0.05. A set of stable variables were defined by LASSO in combination with stability selection. Then coefficients of the selected variables were estimated by fitting generalized linear models using the R function *glm*.

### Network validation using TCGA and GTEx data

Gene expression and gene copy number data of 14 different tumor cohorts (4548 tumor patients in total) from TCGA collected in a previous study(10) were used for validation. We examined the predictive power of our inferred networks on each TCGA cohort by predicting the expression level of each gene for each tumor using the corresponding copy number and gene expression data.

Additionally, in order to validate the applicability of the learnt network to healthy tissues, we further leveraged gene expression data from the Genotype-Tissue Expression (GTEx) Project. Read counts were downloaded from the portal website (version 8), normalized using the R package *DESeq2* and centered gene-wise across tissues.

For each TCGA cohort or GTEx tissue, the expression levels of each gene were predicted using the network and expression quantification of the interacting genes in the same sample. The predicted value was then compared to the observed value, present in the original dataset. The quality of prediction for each TCGA cohort or GTEx tissue was quantified as either the correlation between predicted and observed expression of a gene across all samples or the MSE of prediction of a gene across all samples. A strong positive correlation or high MSE for a gene suggests high predictive power by the network on the respective gene.

### Single-cell test data processing

Human embryonic stem cell differentiation data(12) were downloaded from the Gene Expression Omnibus (GEO, accession number GSE75748) in the format of expected counts. Only snapshot single-cell data were used in this work (file GSE75748_sc_cell_type_ec.csv.gz). The downloaded data were converted to RPM (reads per million). Renal cell carcinoma data(15), in the format of normalized UMI counts, and corresponding metadata, were download via the Single Cell Portal. Data was reduced to cells from patient P915. Cells with library sizes more than 3 median absolute deviations above the median were removed as potential doublets. 2000 cells were randomly selected for further analysis and underwent reversion of the log-transformation. Human embryonic kidney (HEK) cell read and UMI data, sequenced with Smart-seq3 (11), were downloaded from ArrayExpress (accession code E-MTAB-8735). Ensembl IDs were converted to gene symbol and data was normalized for library size (RPM). Oligodendroglioma data(13) were downloaded from GEO (accession number GSE70630) as log_2_(TPM/10+1) and converted back to TPM. Lung Atlas 10X and Smart-seq2 data(14), together with corresponding metadata, were downloaded from Synapse (ID syn21041850). For both sequencing methods, data was restricted to cells from the lung of patient 1. Potential doublets (cells with library sizes above the median) were removed from the 10X data. For this dataset, library sizes above the median were considered potential doublets due to the bi-modal distribution of library sizes below the usual threshold of median+(3*MAD). After doublet removal, 2000 cells were randomly selected for RPM normalization and further analysis for both sequencing methods.

### Dropout imputation

Version 0.0.9 of scImpute(Li and Li 2018) was used for dropout imputation, in “TPM” mode for the oligodendroglioma dataset and “count” mode for all other datasets, without specifying cell type labels. The parameters were left as default, except for drop_thre = 0.3, as the default of 0.5 resulted in no imputations performed. Cell cluster number (Kcluster) was left at the default value of 2 for imputation of all datasets except for the hESC differentiation dataset, where it was set to 6 in order to match the number of cell clusters identified by the authors(Chu et al. 2016), and the Smart-seq3 dataset, where it was set to 1 because only one cell type was sequenced. SAVER 1.1.1 was used with size.factors = 1. SCRABBLE 0.0.1 was run with the parameters suggested by the authors and using by default the average gene expression across cells as the bulk reference. In the case of the hESC differentiation dataset, bulk data from the same study was available, and thus was also used as reference. For all other imputation methods, the data was log2-transformed with a pseudocount of 1. DrImpute 1.0 was run using the default parameters. For dropout imputation by average expression (‘Baseline’), gene expression levels were log2-transformed with a pseudocount of 1 and the average expression of each gene across all cells, excluding zeros, was used for imputation. For network-based imputation, expression values were log2-transformed with a pseudocount of 1 and centered gene-wise across all cells. The original centers were stored for posterior re-conversion. Subsequently, cell-specific deviations of expression levels from those centers were predicted using either equation (5) or the following iterative procedure. During the iteration genes were first predicted using all measured predictors. Subsequently, genes with dropout predictors were re-predicted using the imputed values from the previous iteration. This was repeated for at most 50 iterations. The obtained values were added to the gene-wise centers. We note that the values after imputation cannot be interpreted as TPMs/RPMs, as the sum of the expression levels per sample is no longer guaranteed to be the same across samples. However, one could still perform a new normalization by total signal (sum over all genes) to overcome this issue.

### Masking procedures

In order to compare the imputation error of the tested methods, we randomly masked (set to zero) some of the values for each gene, using two different approaches.

The first approach consisted of setting a fraction of the quantified, uniformly sampled values to zero for each gene (Figure 1 and Supplementary Figs. 4-5) -35% for the hESC differentiation and Smart-seq3 datasets, 10% for the oligodendroglioma and Lung Atlas (FACS-sorted) datasets and 8% for the Renal Cell Carcinoma and Lung Atlas 10X datasets. In case of Supplementary Fig. 6 30% of the cells (not genes) were sampled. This unbiased masking scheme is in agreement with previous work(Talwar et al. 2018). The differing percentages of masked values per gene in each dataset result in a comparable sparsity of the data after masking.

As an alternative masking procedure that represents more closely a downsampling process, we modelled for each gene its probability to be an observed zero in the following way: the fraction of cells where each gene was not captured (zero in the original data) was modelled as a function of its average expression across cells (Supplementary Fig. 8). For this, a cubic spline was used, with knots at each 10% quantile of the average expression levels, excluding the 0% and 100% quantiles. A cubic spline was chosen so that it could properly fit to both UMI-based and non UMI-based datasets. With this model, a ‘dropout probability’ *p* was computed for each gene from its mean expression. The masking procedure then consisted of, for each entry, sampling a Bernoulli distribution with probability of success 1-*p*, where 0 corresponds to a mask (the entry is set to 0) and 1 to leaving the data as it is. Thus, each entry in the data matrix may be masked with a probability *p*, which is gene-specific and based on the observed dropout rates in the dataset at hand.

We observed the same relative performance of the imputation methods under this alternative masking scheme (Supplementary Fig. 9), and for this reason present the results obtained with the first masking approach.

### Imputation performance analysis

Imputation was performed with each of the four tested methods separately and the imputed masked entries were then compared to the original ones. For genes where at least 10 values were imputed (non-zero after imputation, zero after masking) by all methods, the Pearson correlation between original and imputed values across cells was computed for each gene individually.

Additionally, we used the mean of the squared imputation error across all imputations for a given gene:

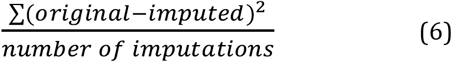

In order to avoid higher errors for more highly expressed genes, we split the genes into expression quartiles when reporting the imputation error.

### Dimensionality reduction and marker detection

Dimensionality reduction on the original hESC differentiation data (Fig. 4B) was performed using ZINBWaVe, t-SNE and UMAP (Supplementary Figs. 12-13). H1 and TB cells in Batch 3 were removed to avoid confounding batch effects and dimensionality reduction was performed for the remaining cells. UMAP was performed on the first 5 principal components obtained from the top 1000 most variable genes in the hESC differentiation data (normalized, log_2_-transformed) before and after imputation (Supplementary Fig. 12) using the *Seurat* R package(Butler et al. 2018). ZINB-WaVe, implemented in the R package *zinbwave*(Risso et al. 2018), was used to extract 2 latent variables from the information contained in the top 1000 genes with highest variance across cells. Batch information and the default intercepts were included in the ZINB-WaVe model, using *epsilon* = 1000 (Supplementary Fig. 13). K-means clustering (k = 6) on the 2 latent variables strongly matched the annotated cell type labels (0.977 accuracy), confirming the reliability of this approach. t-SNE was performed on the normalized and log_2_-transformed data using the *Rtsne* R package with default settings (Supplementary Fig. 13).

Cluster-specific markers were detected from the log_2_-transformed normalized data using Seurat. Detection rate was regressed out using the *ScaleData* function with *vars*.*to*.*regress* = *nGene*. Markers were detected with the *FindAllMarkers* function, using MAST(Finak et al. 2015) test and setting *logfc*.*threshold* and *min*.*pct* to 0, and *min*.*cells*.*gene* to 1.

### GO term enrichment and transcription factor analyses

All GO term enrichment analyses were performed with the *topGO* R package(Alexa et al. 2006). Enrichment in GO biological process terms among cluster-specific markers (Fig. 3C-D) was performed for each cell cluster and (no) imputation method separately, using as foreground the set of significant cluster markers detected by Seurat, with FDR < 0.05 and |logFC| > 0.25, and as background all genes in the Seurat result (both significant and non-significant). The *classic* algorithm was used, in combination with Fisher test, and log_2_ enrichment was quantified as the log_2_ of the ratio between the number of significant and expected genes in each term. Significantly enriched (p-value < 0.001 and log_2_ enrichment > 0.5) GO biological process terms within each set of cluster markers, as detected in the original data (no masking, no imputation), were defined as “high confidence” terms.

For regulatory GO molecular function term enrichment analyses (Fig. 4A), significant (FDR < 0.05 and |logFC| > 0.25) markers uniquely detected without / with each imputation method were combined across all clusters and tested for enrichment in the term “molecular function regulator” against the background of all genes obtained as the result of Seurat (both significant and non-significant). The *classic* algorithm was used, in combination with Fisher test, and log_2_ enrichment was quantified as the log_2_ of the ratio between the number of significant and expected genes in each term.

To identify transcription factors (TFs) among cluster markers exclusively detected using the network-based method, a curated TF list was downloaded from http://www.tfcheckpoint.org/index.php/browse.

### Determination of the optimal imputation method per gene

In order to determine the best performing imputation method for each gene, 70% of the cells in each dataset were used as training data, where a percentage of the expression values were masked, as previously described. In the Smart-seq3 dataset, where only around 100 cells were available, the training was done in 98% of the dataset The remaining cells were used for testing. After masking, each of the tested imputation methods was applied to the training data and the imputed values of masked entries were then compared to the measured values. The Pearson correlation coefficient was computed for each gene with at least 10 imputed (with a non-zero value) masked entries. For each gene, the method leading to the highest correlation coefficient was chosen as optimal. When no decision could be done, the Baseline method was used as a default.

### The ADImpute R package

The ADImpute R package is composed of two main functions, *EvaluateMethods* and *Impute. EvaluateMethods* determines, for each gene, the method resulting in the best imputation performance, in a cross-validation procedure. *Impute* performs dropout imputation according to the choice of method provided by the user. Currently supported methods are scImpute, DrImpute, SAVER, SCRABBLE, the Baseline and Network methods described in this manuscript and an Ensemble method, which takes the results from *EvaluateMethods* to select the imputation results from the gene-specific best method. Additionally, the user can choose to estimate the probability that each dropout value is a true zero, according to the approach used by scImpute, and leave the values unimputed if their probability of being a true zero falls above a user-defined threshold.

### Data access

The data used in this study are publicly available, as described in the Methods section. Human embryonic stem cell differentiation data are available in GEO under the accession number GSE75748. Renal cell carcinoma data are available in the Single Cell Portal. Human embryonic kidney (HEK) cell data are available in ArrayExpress (accession code E-MTAB-8735). Oligodendroglioma data are available in GEO under accession number GSE70630. Lung Atlas 10X and Smart-seq2 data are available in Synapse (ID syn21041850). The ADImpute R package, including the transcriptional regulatory network used in this study, is available via Bioconductor: https://bioconductor.org/packages/release/bioc/html/ADImpute.html.

## Acknowledgements

The results here shown are in part based upon data generated by the TCGA Research Network: http://cancergenome.nih.gov/. The Genotype-Tissue Expression (GTEx) Project was supported by the Common Fund of the Office of the Director of the National Institutes of Health, and by NCI, NHGRI, NHLBI, NIDA, NIMH, and NINDS. The data used for the analyses described in this manuscript were obtained from the GTEx Portal on 12/2019. A.C.L. received support by the Cologne Graduate School of Ageing Research. X.W. received financial support from the National Natural Science Foundation of China (61871463) and Natural Science Foundation of Fujian Province of China (2017J01068).

We gratefully acknowledge help from Dr. Michael Seifert (TU Dresden, Germany) on the construction of the transcriptional regulatory network.

## Author Contributions

AB envisioned the study. XW implemented and performed the network training and testing. ACL implemented and tested the dropout imputation method. All authors contributed to the writing of the manuscript.

