## Supplementary Material for "Regulatory network-based imputation of dropouts in single-cell RNA sequencing data"

**Supplementary Table 1: Characteristics of the test datasets.**

| Authors | Tissue | Sequencing method | # cells | Avg. quantified genes / cell |
|---|---|---|---|---|
| Hagemann-Jensen *et al.* | Human Embryonic Kidney (HEK) cells | Smart-seq3 | 117 | 10,743 (read counts) |
| Hagemann-Jensen *et al.* | Human Embryonic Kidney (HEK) cells | Smart-seq3 | 117 | 9,539 (UMI counts) |
| Chu *et al.* | hESC differentiation | Smart-seq/C1 | 1,018 (snapshot) | 9,617 (snapshot) |
| Tirosh *et al.* | Oligodendroglioma, 6 donors | Smart-seq2 | 4,347 | 5,169 |
| Travaglini *et al.* | Healthy lung, 3 donors' lung and blood | 10X<br>Smart-seq2 | 7,524 (10X, donor 1 lung)<br>3,235 (Smart-seq2, donor 1 lung) | 3,003 (Smart-seq2, donor 1 lung) |
| Travaglini *et al.* | Healthy lung, 3 donors' lung and blood | 10X<br>Smart-seq2 | 7,524 (10X, donor 1 lung)<br>3,235 (Smart-seq2, donor 1 lung) | 2,105 (10X, donor 1 lung) |
| Bi *et al.* | Clear cell renal cell carcinoma, 8 donors | 10X | 6,541 (donor P915) | 1,473 (donor P915) |

**Supplementary Table 2: Spearman rank correlation between original values and imputation results after masking.** Correlation was computed between the vector of original entries and imputations common to all methods.

| | Baseline | DrImpute | SAVER | scImpute | SCRABBLE, bulk | SCRABBLE, sc ref | Network |
|---|---|---|---|---|---|---|---|
| **Smart-seq3 (reads)** | 0.79 | 0.72 | 0.75 | 0.78 | - | 0.72 | 0.79 |
| **Smart-seq3 (UMIs)** | 0.77 | 0.72 | 0.75 | 0.77 | - | 0.70 | 0.77 |
| **hESC differentiation** | 0.62 | 0.62 | 0.43 | 0.61 | | 0.60 | 0.62 |
| **Oligodendroglioma** | 0.65 | 0.41 | 0.29 | 0.43 | - | 0.45 | 0.64 |
| **Lung Atlas (10X)** | 0.66 | 0.60 | 0.49 | 0.47 | - | 0.47 | 0.71 |
| **Lung Atlas (FACS)** | 0.52 | 0.41 | 0.34 | 0.39 | - | 0.41 | 0.53 |
| **ccRCC** | 0.65 | 0.58 | 0.57 | 0.51 | - | 0.48 | 0.72 |

**Supplementary Table 3: Percentage of masked dropouts imputed by each method in the tested datasets.**

| | Baseline | DrImpute | SAVER | scImpute | SCRABBLE, bulk | SCRABBLE, sc ref | Network |
|---|---|---|---|---|---|---|---|
| **Smart-seq3 (reads)** | 81.8% | 79.9% | 81.8% | 80.7% | - | 82.2% | 75.7% |
| **Smart-seq3 (UMIs)** | 80.7% | 78.9% | 80.7% | 80.0% | - | 82.8% | 78.0% |
| **hESC differentiation** | 84.1% | 83.7% | 84.1% | 78.9% | 63.5% | 37.8% | 87.6% |
| **Oligodendroglioma** | 84.8% | 84.1% | 84.8% | 80.6% | - | 24.9% | 85.2% |
| **Lung Atlas (10X)** | 51.1% | 49.3% | 51.1% | 20.0% | - | 18.8% | 68.6% |
| **Lung Atlas (FACS)** | 65.6% | 64.8% | 65.6% | 62.3% | - | 15.6% | 71.2% |
| **ccRCC** | 59.7% | 58.4% | 59.7% | 29.1% | - | 19.5% | 73.7% |

**Supplementary Table 4: Percentage of genes best imputed by each method (lowest MSE) in the seven test datasets restricted to values that could be imputed by all methods.**

| | Baseline | DrImpute | SAVER | scImpute | SCRABBLE, bulk | SCRABBLE, sc ref | Network |
|---|---|---|---|---|---|---|---|
| **Smart-seq3 (reads)** | **47.0%** | 0% | 0% | 9.4% | - | 1.2% | 42.5% |
| **Smart-seq3 (UMIs)** | **48.2%** | 0% | 0% | 5.2% | - | 0.2% | 46.4% |
| **hESC differentiation** | **33.9%** | 0.5% | 0.1% | 21.6% | 4.5% | 5.5% | **33.9%** |
| **Oligodendroglioma** | **47.8%** | 0.2% | 0% | 4.6% | - | 1.9% | 45.5% |
| **Lung Atlas (10X)** | 16.7% | 0.5% | 0% | 0% | - | 0.4% | **82.5%** |
| **Lung Atlas (FACS)** | 35.6% | 0.1% | 0% | 2.2% | - | 3.3% | **58.8%** |
| **ccRCC** | 12.6% | 0.4% | 0% | 0% | - | 0.4% | **86.6%** |

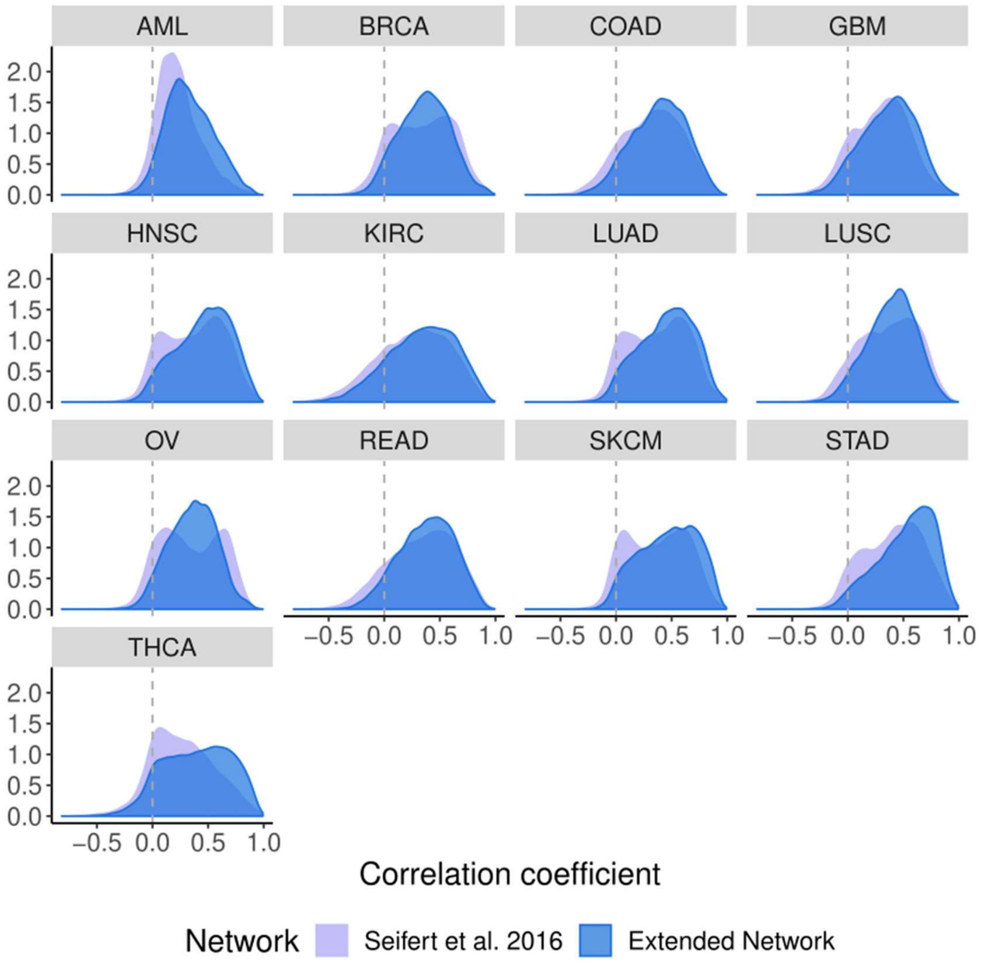


**Supplementary Figure 1: Correlation between network predictions (using the model from Seifert et al.[9] (light purple) and the improved network described here (blue)) and measured gene expression in diverse TCGA datasets.** For each gene its expression was predicted in a given tumor sample using the measured expression values of all detected predictors in the model. Subsequently, observed and predicted values were correlated across all samples from one cohort. The plots show the distributions of Pearson's correlation scores across all genes common between the network model and the respective TCGA dataset. Although there is variation with respect to how well genes in different tumor entities can be predicted, the distributions are always strongly skewed in favour of positive correlations. This trend is enhanced with the new model presented here. AML - Acute Myeloid Leukemia; BRCA - Breast Invasive Carcinoma; COAD - Colon Adenocarcinoma; GBM - Glioblastoma Multiforme; HNSC - Head and Neck Squamous Cell Carcinoma; KIRC - Kidney Renal Clear Cell Carcinoma; LUAD - Lung Adenocarcinoma; LUSC - Lung Squamous Cell Carcinoma; OV - Ovarian Serous Cystadenocarcinoma; READ - Rectum Adenocarcinoma; SKCM - Skin Cutaneous Melanoma; STAD - Stomach Adenocarcinoma; THCA - Thyroid Carcinoma.

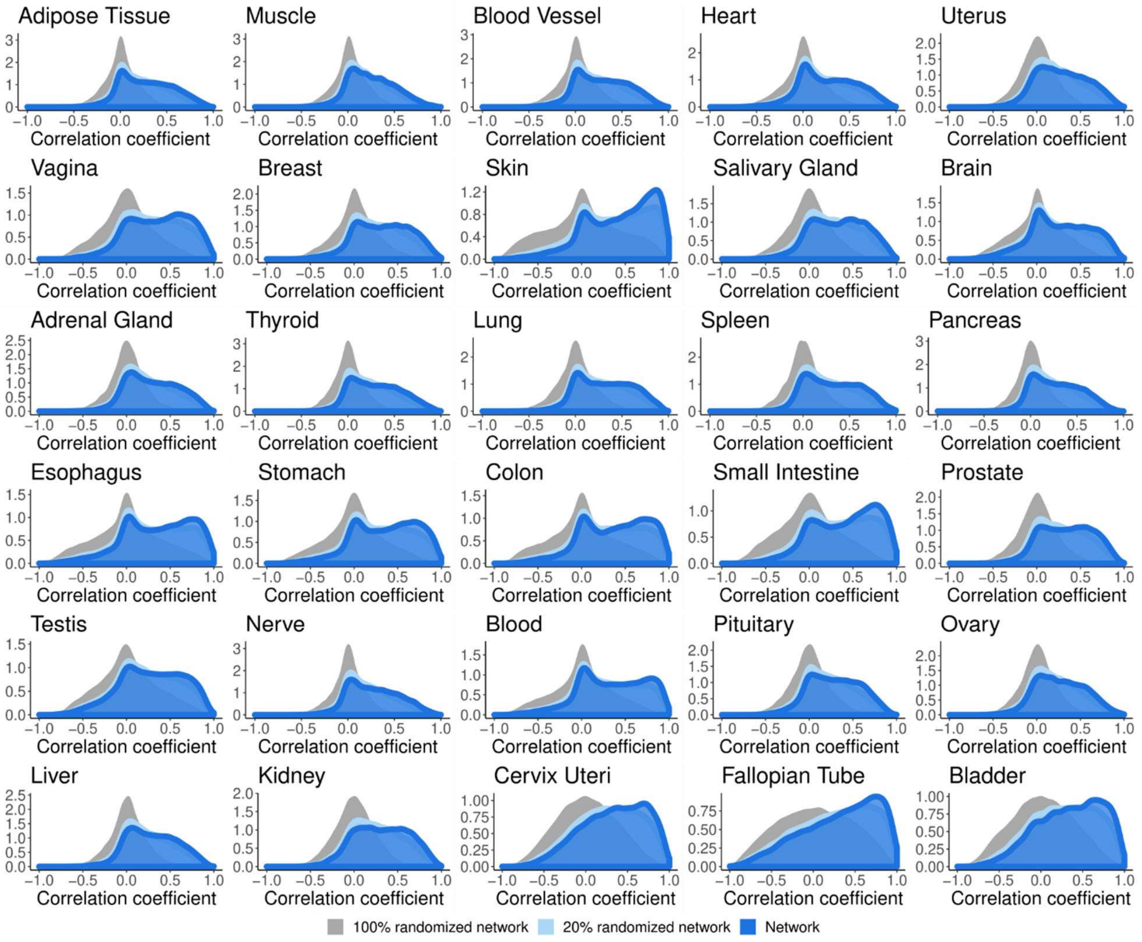


**Supplementary Figure 2: Correlation between network predictions (using the network described here – blue - , a partially randomized network - light blue - and a fully randomized network – grey) and measured gene expression in diverse healthy tissues from the GTEx consortium.** For each gene its expression was predicted in a given healthy tissue sample using the measured expression values of all detected predictors in the model. Subsequently, observed and predicted values were correlated across all samples from one tissue. The plots show the distributions of Pearson's correlation scores across all genes common between the network model and the GTEx data.

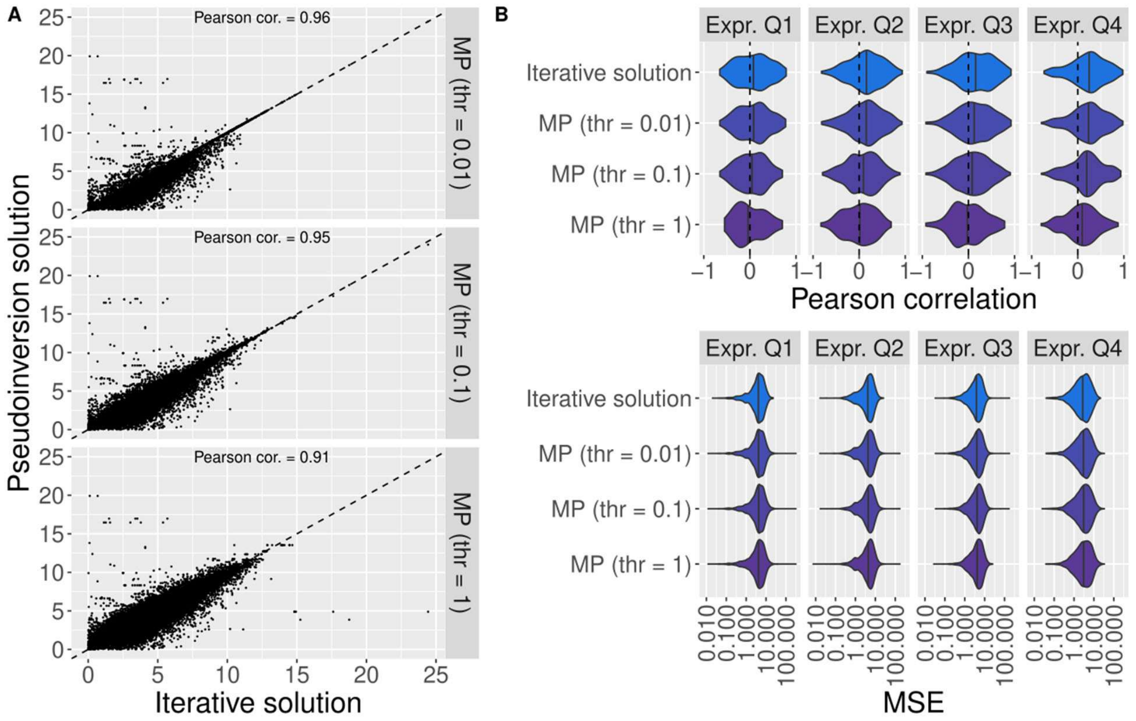


**Supplementary Figure 3: Comparison between results of the network-based imputation using the iterative approach and the Moore-Penrose pseudoinversion (MP) with varying tolerance thresholds in a random subset of 20 cells from the hESC differentiation dataset.** A) Correlation between the results of the iterative approach (x axis) and the Moore-Penrose pseudoinversion (y-axis), across the 20 random cells. B) Imputation performance per gene using the iterative approach and MP with different tolerance thresholds (Pearson correlation, top, and MSE, bottom), separated by expression quartile on the masked data. The higher the tolerance threshold, the fewer singular values are used for the pseudoinversion. Results were limited to imputations performed by all methods.

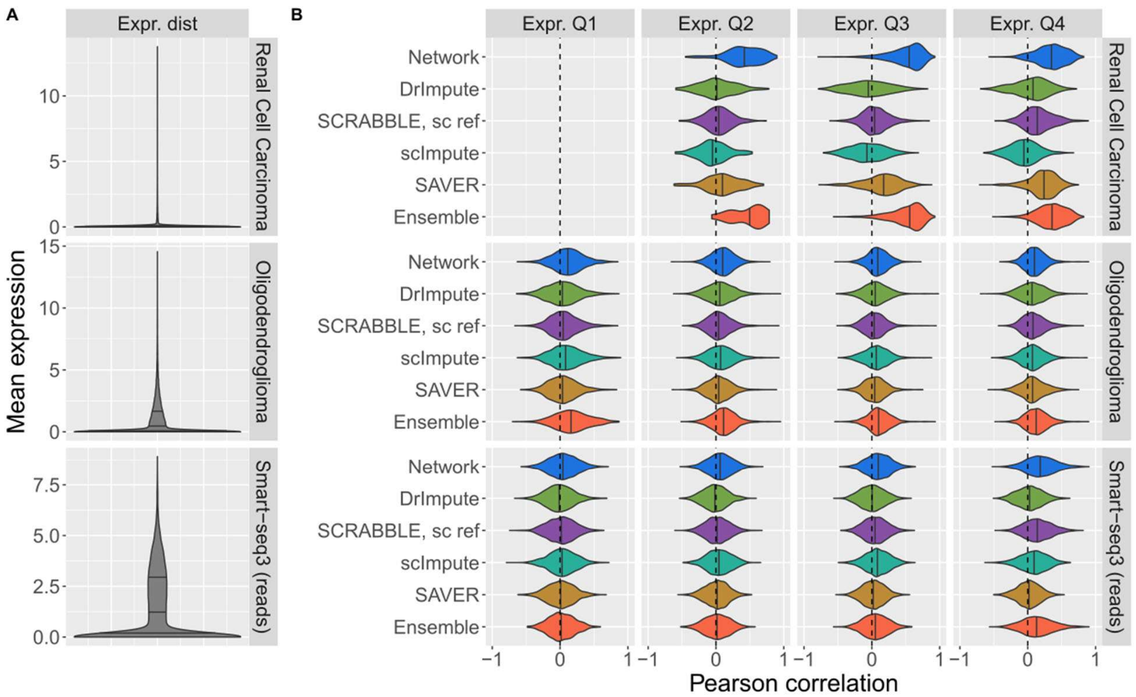


**Supplementary Figure 4: Imputation performance of Network (blue), DrImpute (green), SAVER (yellow), scImpute (turquoise), SCRABBLE (with bulk or single-cell data as reference; purple), Network (blue) and Ensemble (orange), stratified by expression quartiles, on additional datasets. A)** Distribution of average expression levels in each dataset. Quartiles are represented by vertical lines. **B)** Pearson correlation coefficient, for each gene, between the imputation by the specified method and the original values before masking. Only values that could be imputed by all methods were used for correlation computation. Expression quartiles are determined for each dataset separately, on the masked data.

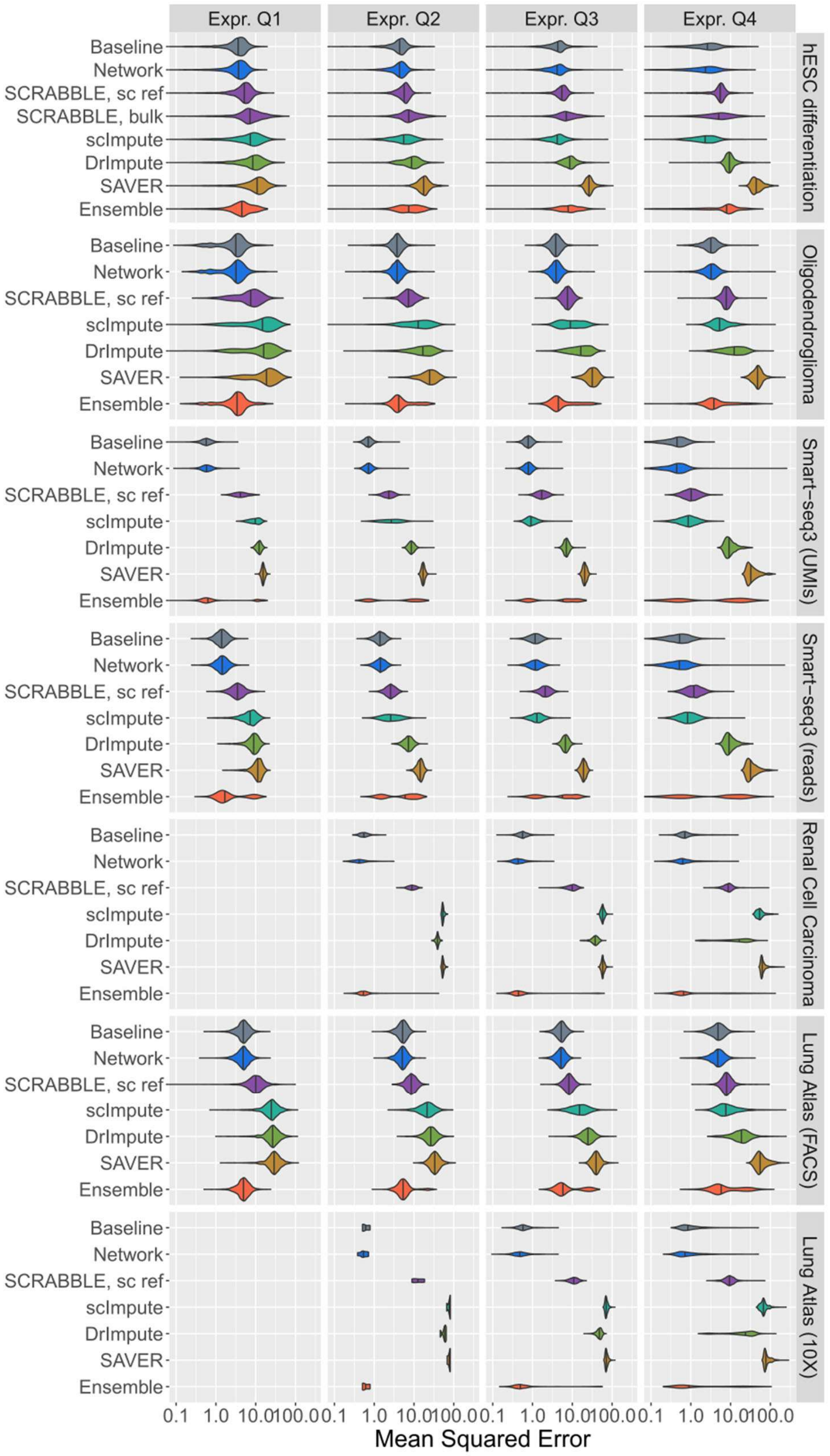


**Supplementary Figure 5: Imputation error of Baseline (slate grey), DrImpute (green), SAVER (yellow), scImpute (turquoise), SCRABBLE (with bulk or single-cell data as reference; purple) and Network (blue), stratified by expression quartiles.** Only values that could be imputed by all methods were used for MSE computation. Expression quartiles are determined for each dataset separately, on the masked data. The x axis is presented log-transformed and was cropped at 0.1 to exclude the low-MSE tail from visualization and facilitate result comparison.

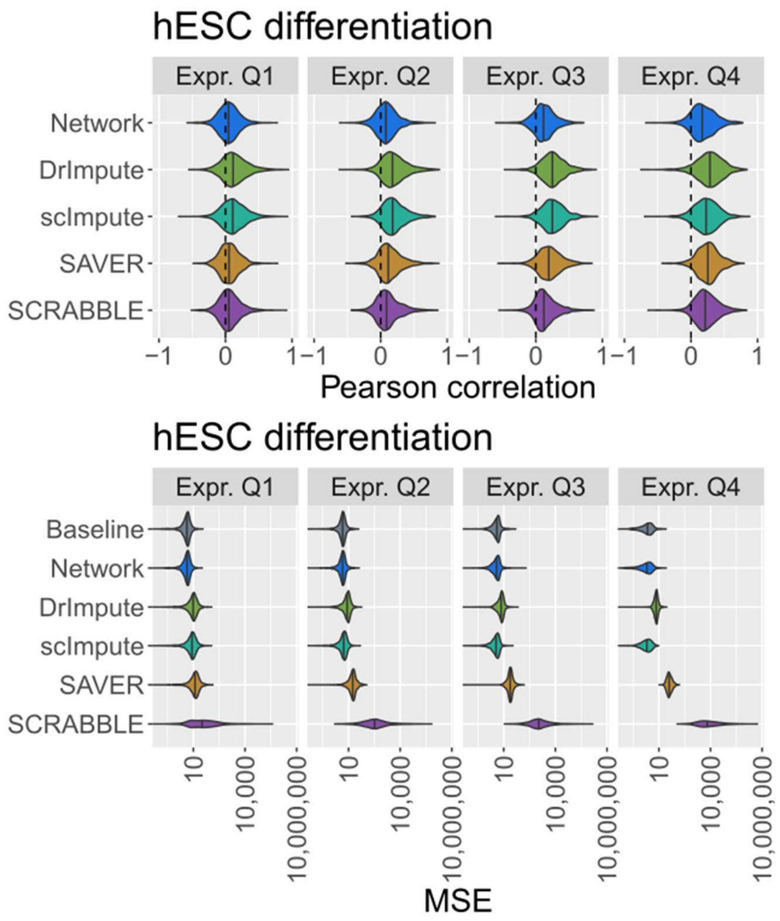


**Supplementary Figure 6: Imputation performance of Baseline (slate grey), Network (blue), DrImpute (green), scImpute (turquoise), SAVER (yellow) and SCRABBLE (purple) methods in the hESC differentiation dataset, upon cellwise masking**. Correlation coefficient between imputed and original values (top) and MSE of imputation (bottom), upon random masking of 30% of the quantified genes in each cell in the hESC differentiation dataset. The Baseline method computes the average expression of a gene across all cells and it is not using the information of any other genes. Hence, one would assume that its error should be independent of the number of missing genes per cell. This is however not the case due to a biased gene sampling: cells with few detected genes will preferentially report values for highly expressed genes, whereas cells with many detected genes will represent a less biased sample of the whole transcriptome. This is affecting the performance, which thus is slightly dependent on the number of missing genes per cell. Values were restricted to imputations performed by all tested methods.

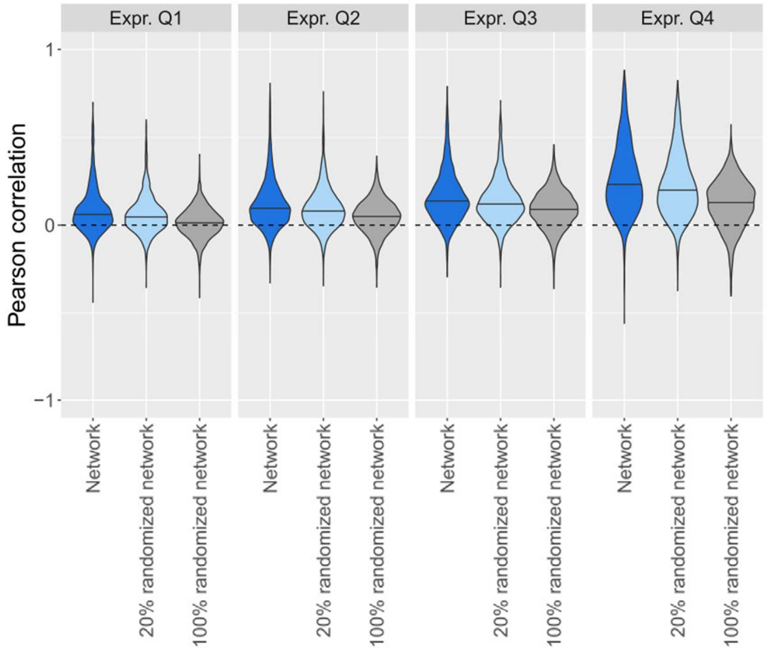


**Supplementary Figure 7: Imputation performance of the Network method upon randomization in the hESC differentiation dataset.** Dropout imputation was performed using the network described here (blue), a partially randomized network (light blue) and a fully randomized network (grey).

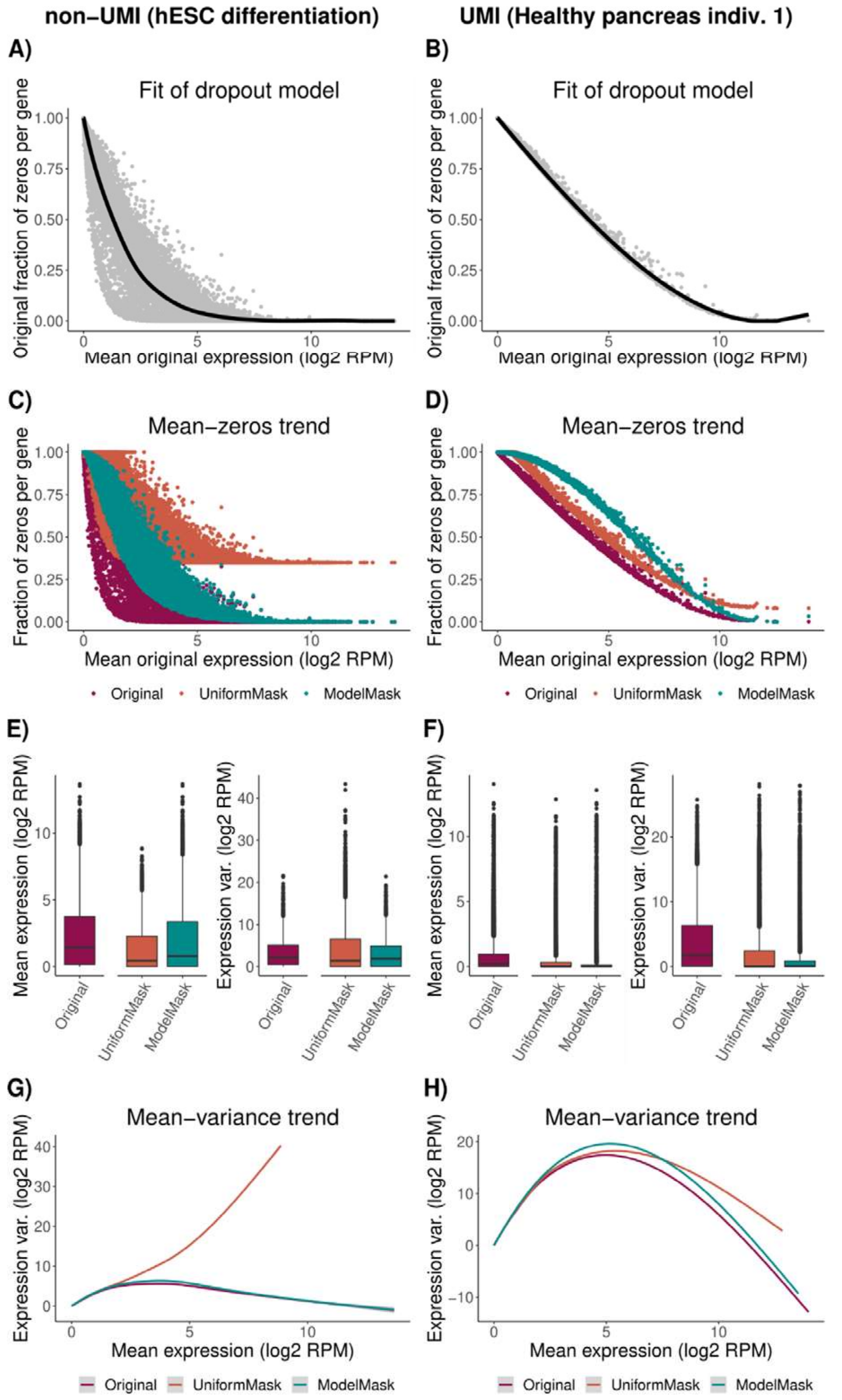


**Supplementary Figure 8: Comparison of the two masking procedures employed in this work: a uniform masking procedure which sets to zero the same number of entries per gene (UniformMask) and a model-based procedure which sets entries to zero with a gene-specific probability obtained from the data.** Comparisons were done on representative datasets of non-UMI data (**A)**, **C)**, **E)**, **G)**) and UMI data (**B)**, **D)**, **F)**, **H)**). **A)** and **B)** Fit of the spline model (Methods) to the original data. **C)** and **D)** Fraction of zeros in the data before (Original) and after (UniformMask, ModelMask) masking, compared to original average gene expression,. **E)** and **F)** Distribution of mean expression and expression variance before and after masking, compared to original average gene expression. **G)** and **H)** Mean-variance trend before and after masking.

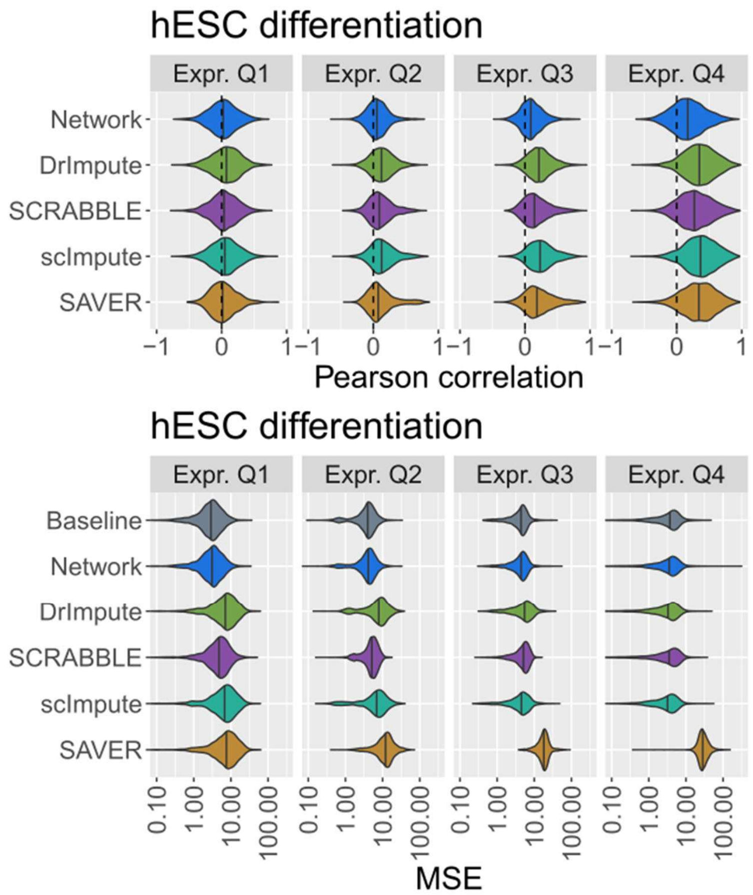


**Supplementary Figure 9: Imputation performance of Baseline (slate grey), DrImpute (green), SAVER (yellow), scImpute (turquoise), SCRABBLE (purple) and Network (blue) methods in the hESC differentiation dataset, using a gene-specific masking procedure (Methods).** Correlation coefficient between imputed and original values (top) and MSE of imputation (bottom), upon random masking of 30% of the quantified genes in each cell in the hESC differentiation dataset. Only values that could be imputed by all methods were used for performance analysis. Expression quartiles are determined on the masked data. The MSE axis is presented log-transformed and was cropped at 0.1 to exclude the low-MSE tail from visualization and facilitate result comparison.

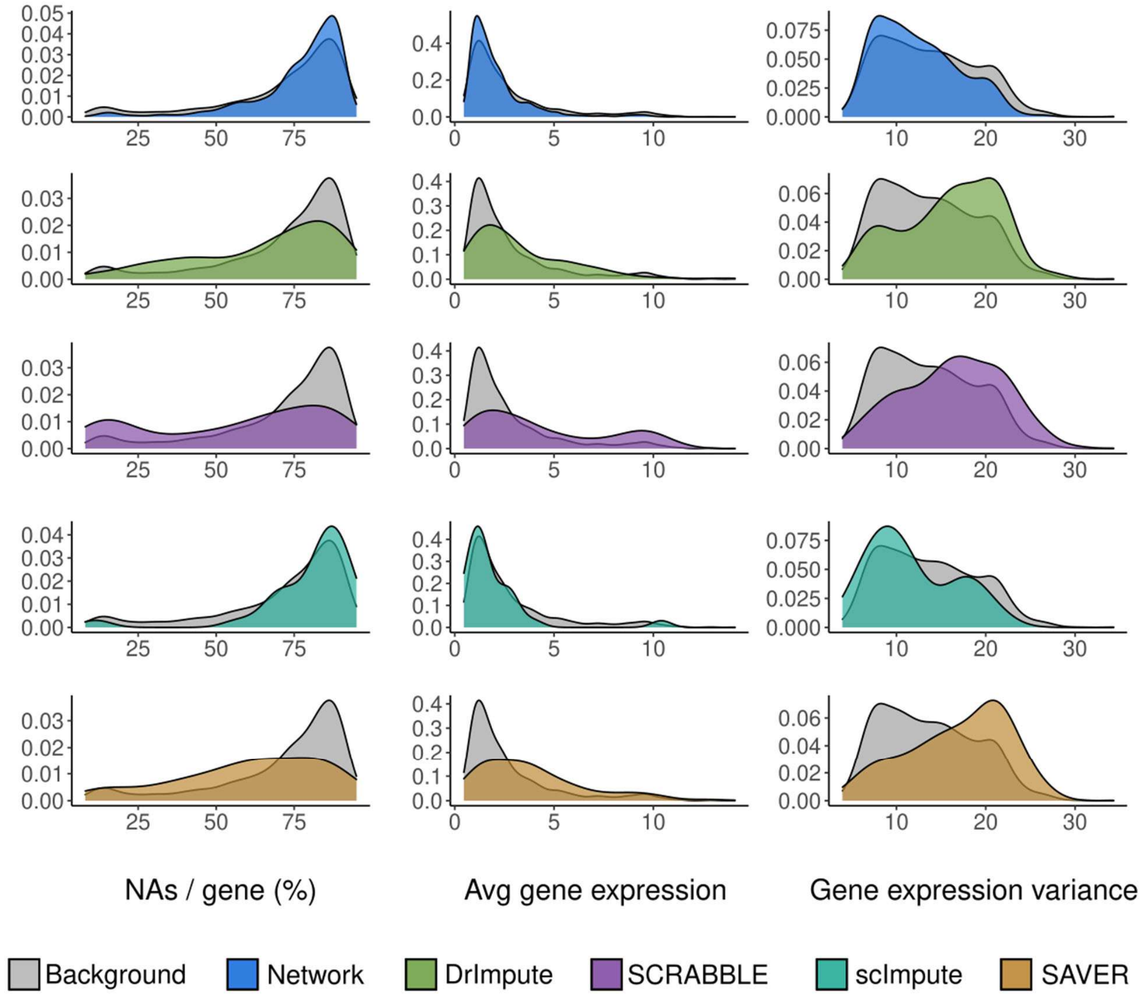


**Supplementary Figure 10: Characterization of the genes best predicted by DrImpute (green), SAVER (yellow), scImpute (turquoise), SCRABBLE (single-cell reference; purple) or Network (blue) methods in the Lung Atlas 10X dataset.** Distribution of missing values per gene, average expression levels and variance of the genes best predicted by Baseline, scImpute and Network methods, compared against all tested genes (background).

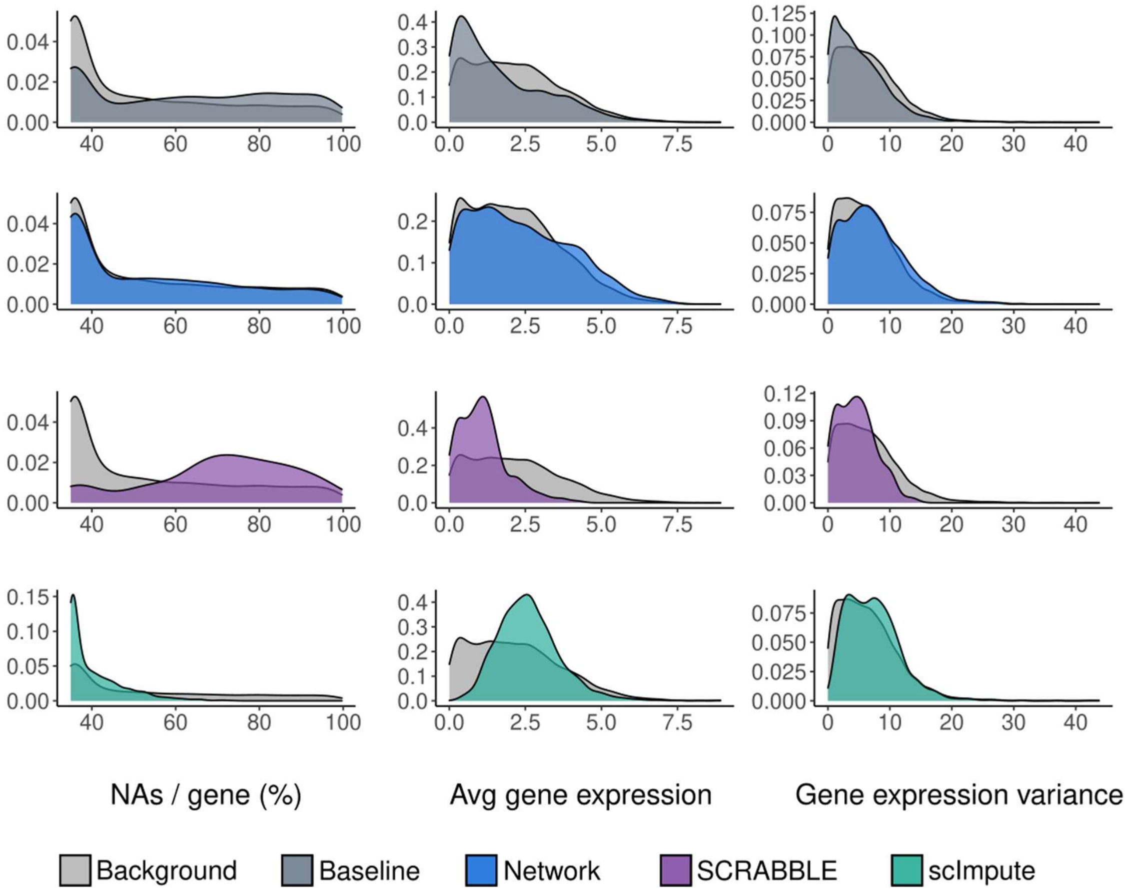


**Supplementary Figure 11: Characterization of the genes best predicted by Baseline (slate grey), scImpute (turquoise), SCRABBLE (purple) or Network (blue) methods in the hESC differentiation dataset, using imputation error to quantify performance.** Distribution of missing values per gene, average expression levels and variance of the genes best predicted by Baseline, scImpute and Network methods, compared against all tested genes (background). Some distributions are not shown due to the low number of best imputed genes.

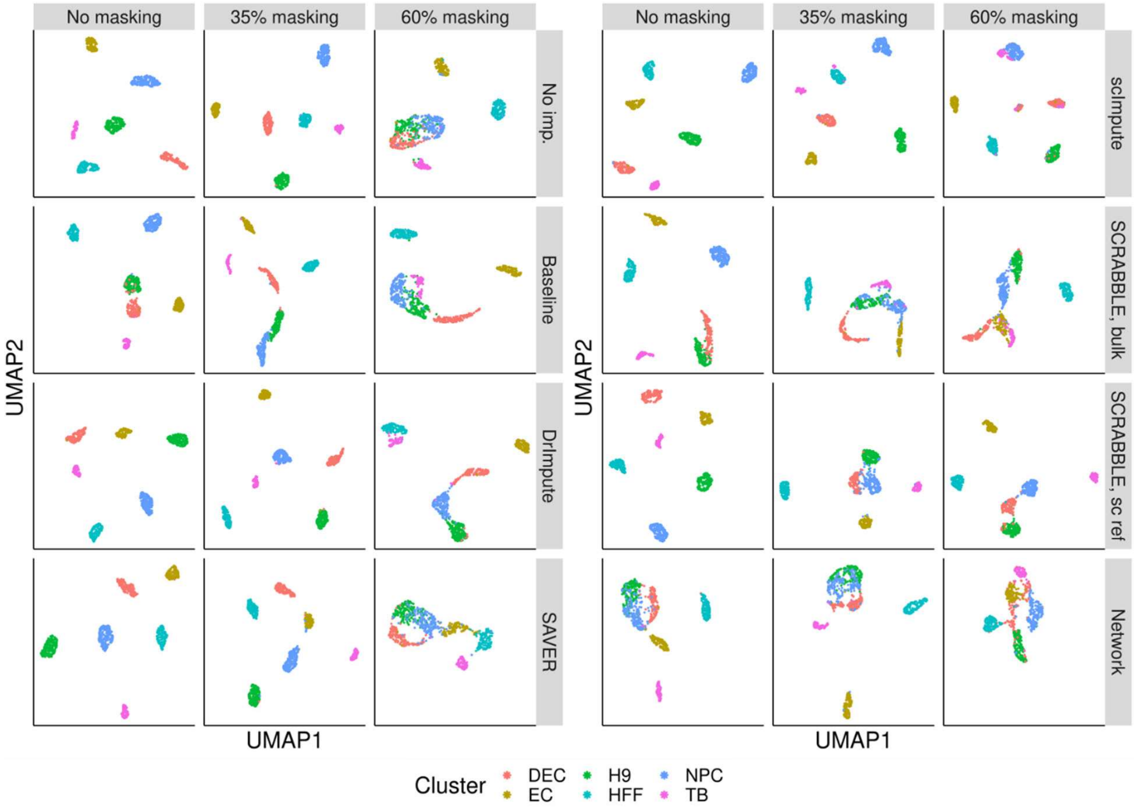


**Supplementary Figure 12: Effect of imputation with Baseline, scImpute, SAVER and Network methods on UMAP plots.** Data was subject to: no masking (left column), relaxed masking (35% of quantified entries per gene were set to zero, middle column), stringent masking (60% of quantified entries set to zero, right column). The plot in the upper left reflects the clustering on the original, unchanged data. Imputation was performed for actually missing values in the original data (all columns) and on masked values (columns 2 & 3). Colors represent cell type label annotations from the original publication. DEC: definitive endoderm cells; EC: endothelial cells; H9: undifferentiated human embryonic stem cells; HFF: human foreskin fibroblasts; NPC: neural progenitor cells; TB: trophoblast-like cells.

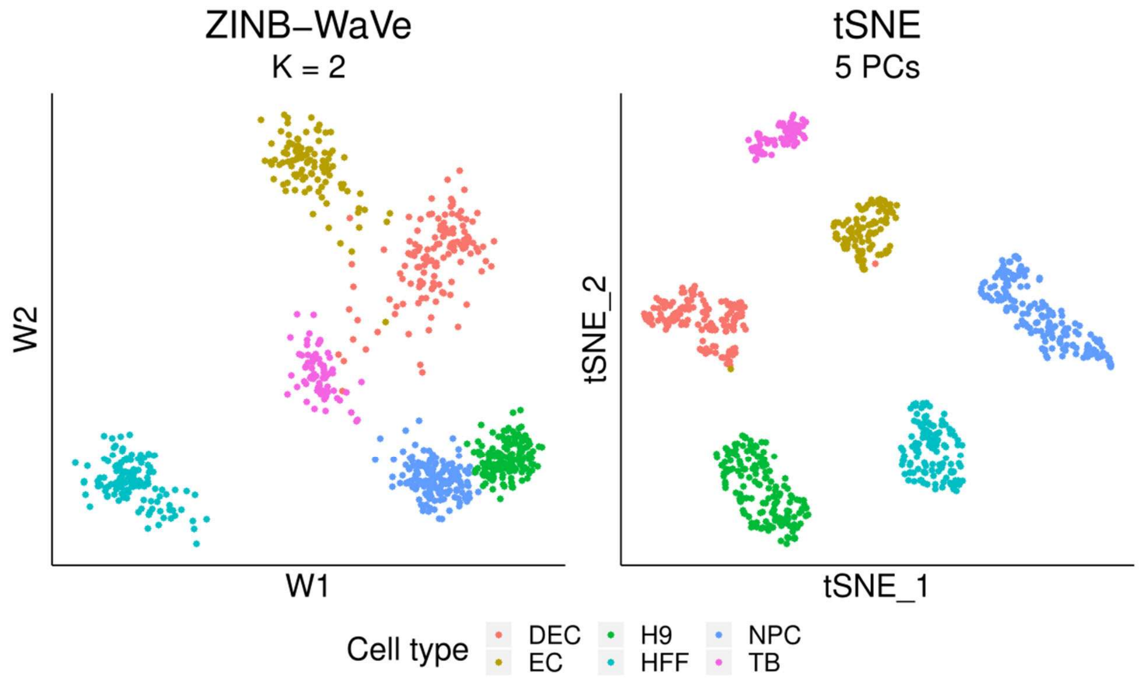


**Supplementary Figure 13: Dimensionality reduction results with different techniques: ZINBWaVe and t-SNE on the hESC dataset.** Data was not subject to any masking. Colors represent cell type label annotations from the original publication. DEC: definitive endoderm cells; EC: endothelial cells; H9: undifferentiated human embryonic stem cells; HFF: human foreskin fibroblasts; NPC: neural progenitor cells; TB: trophoblast-like cells.

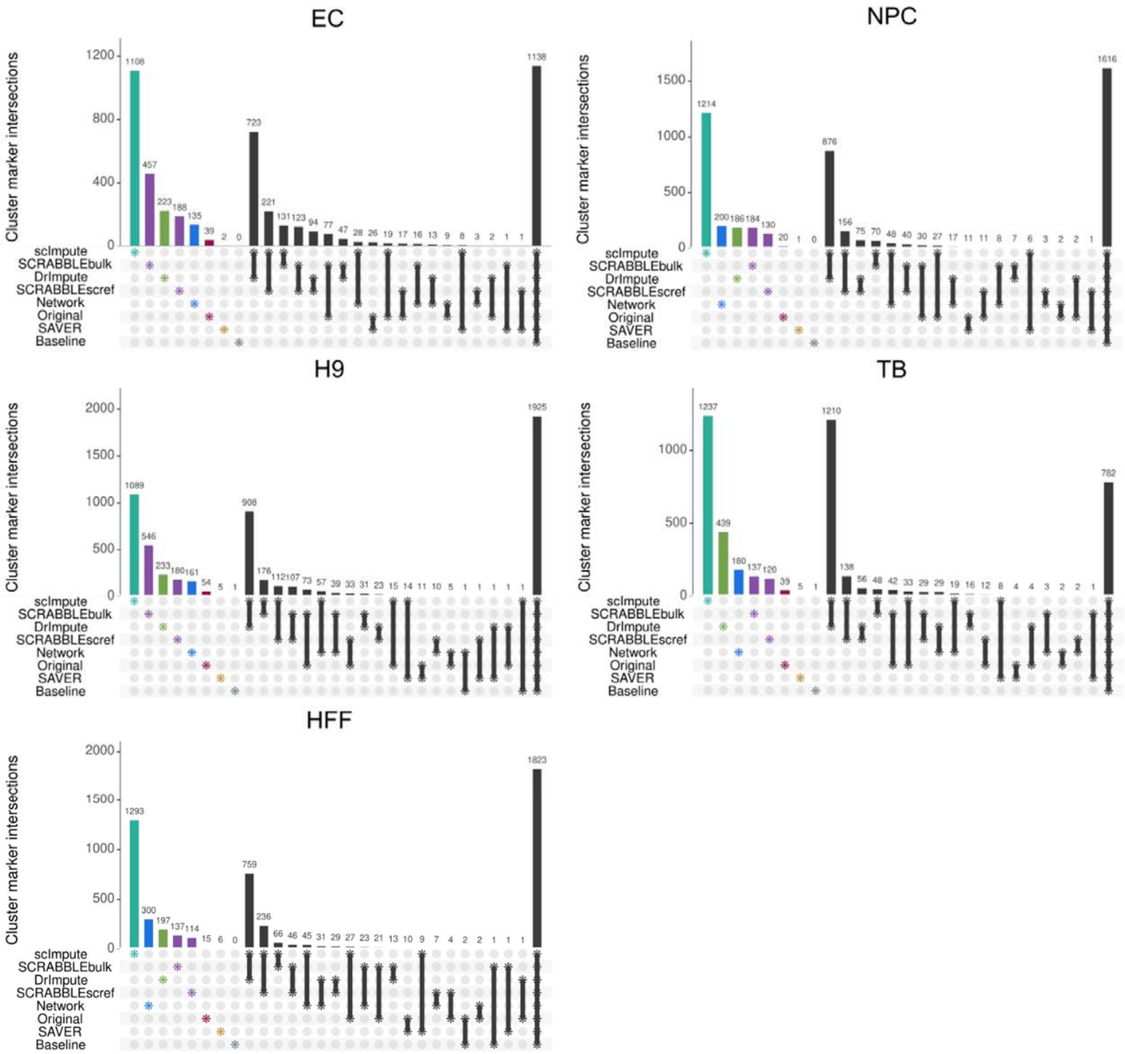


**Supplementary Figure 14: Overlap between significant (FDR < 0.05, $|\log_2 FC| > 0.25$) cell type markers detected with no dropout imputation (Original) and using the tested imputation methods.** DEC: definitive endoderm cells; EC: endothelial cells; H9: undifferentiated human embryonic stem cells; HFF: human foreskin fibroblasts; NPC: neural progenitor cells; TB: trophoblast-like cells.

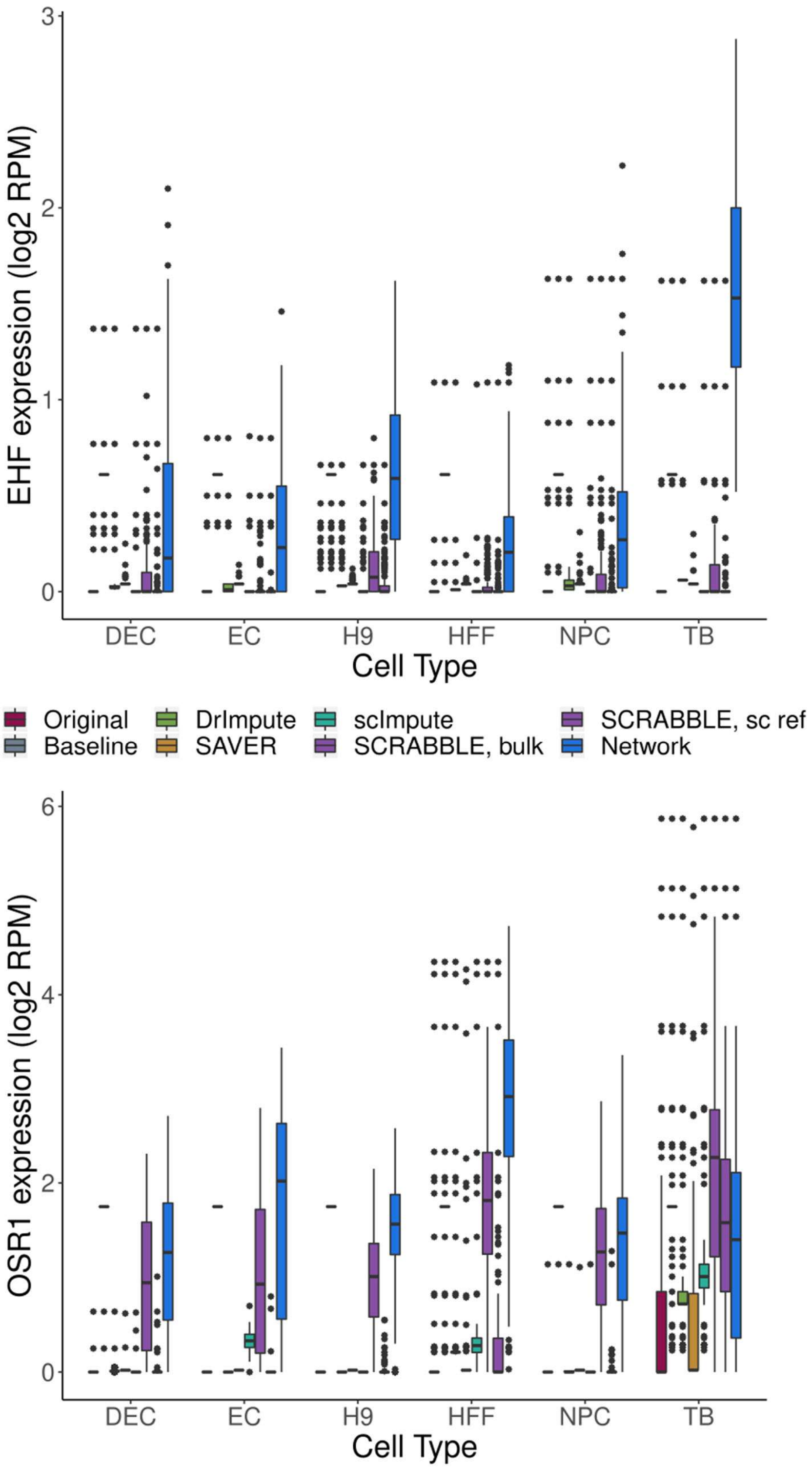


**Supplementary Figure 15: EHF and OSR1 expression levels before and after dropout imputation, across cell types.** DEC: definitive endoderm cells; EC: endothelial cells; H9: undifferentiated human embryonic stem cells; HFF: human foreskin fibroblasts; NPC: neural progenitor cells; TB: trophoblast-like cells.

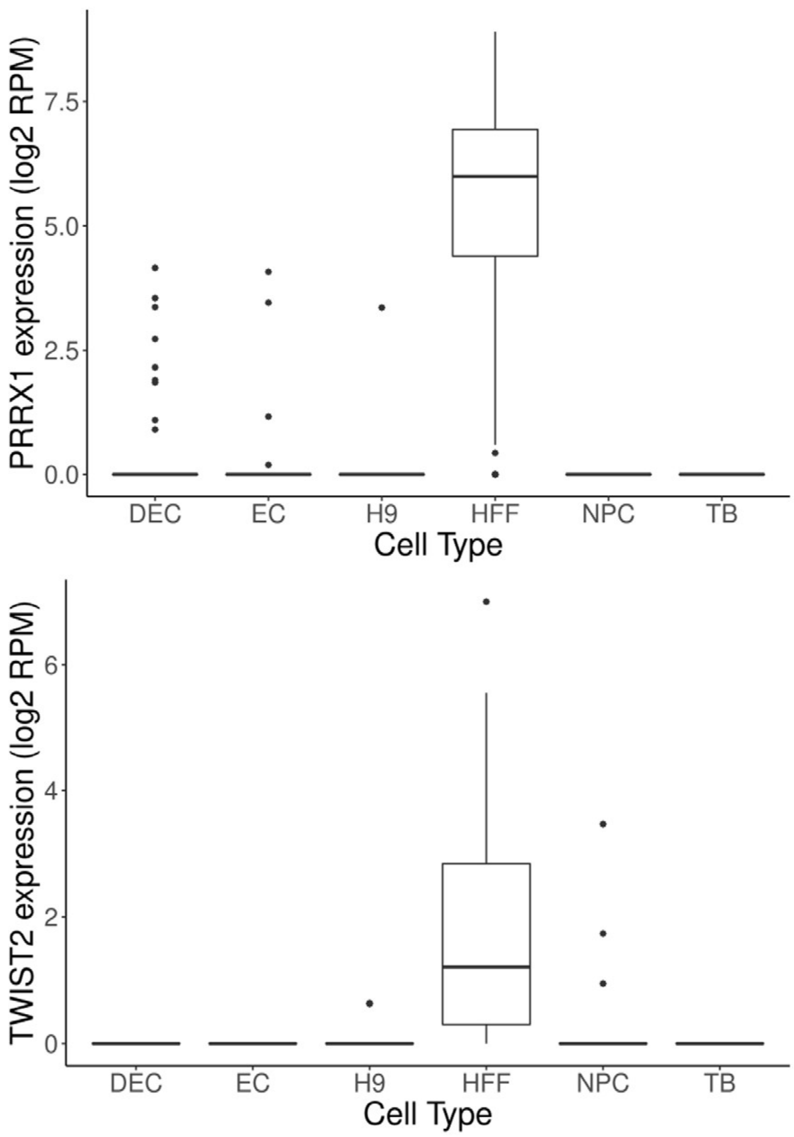


**Supplementary Figure 16: PRRX1 and TWIST2 expression levels before dropout imputation, across cell types.** DEC: definitive endoderm cells; EC: endothelial cells; H9: undifferentiated human embryonic stem cells; HFF: human foreskin fibroblasts; NPC: neural progenitor cells; TB: trophoblast-like cells.
